# c-Src-dependent phosphorylation of Mfn2 regulates endoplasmic reticulum-mitochondria tethering

**DOI:** 10.1101/2022.02.21.481295

**Authors:** Peng Zhang, Kara Ford, Jae Hwi Sung, Yuta Suzuki, Maria Landherr, Jacob Moeller, Isabel Chaput, Iuliia Polina, Madeline Kelly, Bridget Nieto, Toshiaki Tachibana, Yoichiro Kusakari, Michael W. Cypress, Kamelia Drenkova, Stephanie M. Adaniya, Jyotsna Mishra, Ulrike Mende, Bong Sook Jhun, Jin O-Uchi

## Abstract

Contact sites between the mitochondria and endoplasmic reticulum (ER) regulate the exchange of lipids, Ca^2+^, and reactive oxygen species (ROS) across the two organelles. Mitofusin 2 (Mfn2) has been identified as one of the major components tethering these two organelles together. Several post-translational modifications (PTMs) of Mfn2 have been shown to modulate canonical (i.e., mitochondrial fusion) and non-canonical Mfn2 functions, such as mitophagy and activation of ER stress signaling. However, there is little information about whether any PTMs can regulate mitochondrial and ER tethering. Basal tyrosine phosphorylation of Mfn2 was detected by mass spectroscopy, but it is unknown whether Mfn2 is a substrate of mitochondria-localized tyrosine kinases. Here, we show that mitochondria-localized c-Src can phosphorylate the C-terminal tail of Mfn2, which decreases the distance between the mitochondria and ER and facilitates Ca^2+^ transfer from the ER to mitochondria, followed by changes in ROS generation and mitochondrial bioenergetics. Our findings suggest that tyrosine phosphorylation of Mfn2 may uniquely work to fine-tune ER-mitochondrial Ca^2+^ transport under physiological and pathological conditions.

## Introduction

Mitochondria are cellular organelles that play multiple roles in the life and death of a cell, including ATP generation, reactive oxygen species (ROS) generation, apoptosis, and handling of the universal second messenger calcium (Ca^2+^) (Rossmann, Dubois et al. 2021, Murphy, Ardehali et al. 2016). Because mitochondria are semi-autonomous organelles with their own unique DNA, proteins encoded by mitochondrial and nuclear DNA are integrated into the mitochondria to maintain and regulate mitochondrial function (Calvo, Mootha 2010, Rossmann, Dubois et al. 2021). Moreover, mitochondrial function may also be modulated through physical interactions with other organelles, such as the endoplasmic/sarcoplasmic reticulum (ER/SR), peroxisomes, and nucleus (Xia, Zhang et al. 2019, Rossmann, Dubois et al. 2021).

Among these interactions between mitochondria and other organelles, the structural and functional importance of ER/SR and mitochondrial membranes, termed mitochondria-associated membranes (MAM), have been well recognized (Barazzuol, Giamogante et al. 2021, Perrone, Caroccia et al. 2020). MAM is involved in regulating lipid, Ca^2+^, and ROS exchange across these two organelles, which is critical for the maintenance of mitochondrial bioenergetics (Barazzuol, Giamogante et al. 2021, Perrone, Caroccia et al. 2020). Recent studies report that one of the major components of the tethering structure between the two organelles is mitofusin 2 (Mfn2), a dynamin-like GTPase associated with the fusion of the outer mitochondrial membrane (OMM) (Chandhok, Lazarou et al. 2018, Adaniya, O-Uchi et al. 2019). Indeed, Mfn2 ablation inhibits both its canonical function and mitochondrial fusion and also increases the distance between the ER and mitochondria, which decreases agonist-evoked Ca^2+^ transfer from the ER to mitochondria and alters bioenergetics (de Brito, Scorrano 2008, Naon, Zaninello et al. 2016). Based on knockout studies of Mfn2, it has been proposed that Mfn2 also serves as a key regulator of mitophagy and ER stress signaling, which are currently recognized as the non-canonical roles of Mfn2 in addition to ER-mitochondria (ER-Mito) tethering (Ngoh, Papanicolaou et al. 2012, Munoz, Ivanova et al. 2013, Sebastian, Hernandez-Alvarez et al. 2012).

Several post-translational modifications (PTMs), including serine/threonine phosphorylation and ubiquitination, of Mfn2 have been identified that alter the fusion and mitophagy activity (Adaniya, O-Uchi et al. 2019), but it is still unclear whether there are PTMs that regulate its tethering function. Data from mass spectroscopy reveals that there is basal tyrosine phosphorylation (P-Tyr) of Mfn2 (see online database PhosphoSitePlus) (Hornbeck, Zhang et al. 2015). However, the specific signaling pathways that regulate P-Tyr levels of Mfn2 are completely unknown. Non-receptor-type protein tyrosine kinases (PTKs), including multiple Src family kinases (SFKs) and their negative regulator C-terminal Src kinase (CSK), have been observed in the mitochondria as well as the cytoplasm (Miyazaki, Tanaka et al. 2006, Miyazaki, Neff et al. 2003, Tibaldi, Brunati et al. 2008). Among the eight members of the SFK family (Lyn, Hck, Lck, Blk, c-Src, Fyn, Yes, and Fgr) (Okada 2012), four have been found in the mitochondria. Specifically, c-Src, Fyn, Lyn, and Fgr are detectable in the mitochondria by immunoblotting of the crude mitochondrial proteins and by immunogold-labeled transmitted electron microscopy (TEM) (Miyazaki, Tanaka et al. 2006, Miyazaki, Neff et al. 2003, Tibaldi, Brunati et al. 2008).

In this study, we show that Mfn2 is a novel substrate of the main SKF member c-Src and the phosphorylation of Mfn2 results in decreased distance between the OMM and ER at contact sites, which leads to increased Ca^2+^ transfer from ER to mitochondria and mitochondrial ROS (mROS) generation. This study provides a potential candidate for the cellular signaling pathway responsible for fine-tuning the ER-Mito distance via PTM of Mfn2.

## Results

### c-Src is capable of phosphorylating Mfn2

To test whether Mfn2 has the potential to be tyrosine phosphorylated (P-Tyr) by SFKs, we genetically increased SFK activity in HEK293T cells. First, we confirmed that SFK members c-Src, Fyn, and Lyn were detectable in the mitochondrial fraction of HEK293T cells (**Fig. 1A**), similar to a previous report from rat brain mitochondria (Tibaldi, Brunati et al. 2008). In contrast, Fgr, which was shown as another mitochondria-localized SFK in other cell lines (Tibaldi, Brunati et al. 2008), was not observed in this cell line (**Fig. EV1**). Next, to activate SFKs, we generated a stable knock-down (KD) of CSK, a negative regulator of SFKs, by stably overexpressing shRNA targeting CSK or the empty vector PLKO.1 as a control (**Fig. 1B** and **Appendix Fig. S1**). The expression levels of SFKs were not altered by CSK knockdown (CSK-KD), relative to controls transfected with the empty vector PLKO.1 (**Fig. EV1**). To assess the activity of SFKs, total SFK autophosphorylation levels were detected by a specific antibody raised against a synthetic phosphopeptide corresponding to residues surrounding Tyr^419^ of human c-Src, which can cross-react with the autophosphorylation sites of all other SFKs (Du, Wang et al. 2020). In whole cell lysates, we found that SFK activity increased in CSK-KD cells compared to control (**Fig.1B and Appendix Fig. S1**). Importantly, CSK-KD preferentially activates SFK activity in the mitochondria compared to the cytosol (**Fig. 1B**). Next, to test whether CSK-KD can modulate the P-Tyr levels of Mfn2, HA-tagged human Mfn2 was transiently expressed in CSK-KD cells and control cells, and P-Tyr levels in immunoprecipitated Mfn2 were detected by a general P-Tyr antibody. We found that CSK-KD significantly increased P-Tyr levels of Mfn2 (**Fig. 1C and D**). To further investigate which SFK member(s) found in the mitochondrial protein fraction (Miyazaki, Tanaka et al. 2006, Miyazaki, Neff et al. 2003, Tibaldi, Brunati et al. 2008) can phosphorylate Mfn2, we first overexpressed wild-type c-Src (c-Src-WT), a main SFK member, and its kinase-inactive dominant-negative mutant c-Src-K295R (c-Src-DN) in HEK293T cells stably overexpressing human Mfn2-HA and assessed P-Tyr levels in immunoprecipitated Mfn2 (**Fig. 1E and F**). We found that P-Tyr levels of Mfn2 significantly increased after c-Src-WT overexpression (but not in c-Src-DN overexpression) compared to control cells (**Fig. 1F**). Expression of constitutively active c-Src (c-Src-Y416F, termed c-Src-CA (Nagashima, Maruyama et al. 2008))) was also able to phosphorylate Mfn2 assessed by the increased intensity in a slower migrating band using phosphate-affinity electrophoresis technique (Phos-tag SDS-PAGE (Kinoshita, Kinoshita-Kikuta et al. 2006))) (**Fig. 1G**). Using the same experimental system, we also confirmed that overexpressed c-Src was capable of phosphorylating endogenous Mfn2 in primary cells (i.e., adult human cardiac fibroblasts [HCFs] from a healthy donor (Rizvi, DeFranco et al. 2016))) (**Fig. 1H**). In addition, stimulation of fibroblast growth factor (FGF) receptors, which can transiently activate c-Src (peak at 30 min), also increased Mfn2 phosphorylation in HCFs (**Fig. 1I**). Next, we explored the possibility of Mfn2 phosphorylation by other mitochondrial localized SFK members (Fyn, Lyn, and Fgr, see **Fig. 1A**). Although the reported substrates of CSK are only in the SFK family, we also tested whether CSK can directly phosphorylate Mfn2 because CSK is also localized in the mitochondria (see **Fig. 1A and B**). Overexpression of these PTKs was not able to increase P-Tyr of Mfn2 (**Fig. 1J and K**). Lastly, we used a series of proteinase K digestions on isolated mitochondria from HEK293T cells to differentially permeabilize the OMM and inner mitochondrial membranes (IMM) and assess the localization of c-Src. The mitochondria digestion assay revealed that c-Src is abundant in the OMM, where Mfn2 is located, rather than in intra-mitochondrial space (**Fig. 1L and M**), indicating that c-Src located at the OMM is capable of phosphorylating Mfn2.

**Fig. 1.**
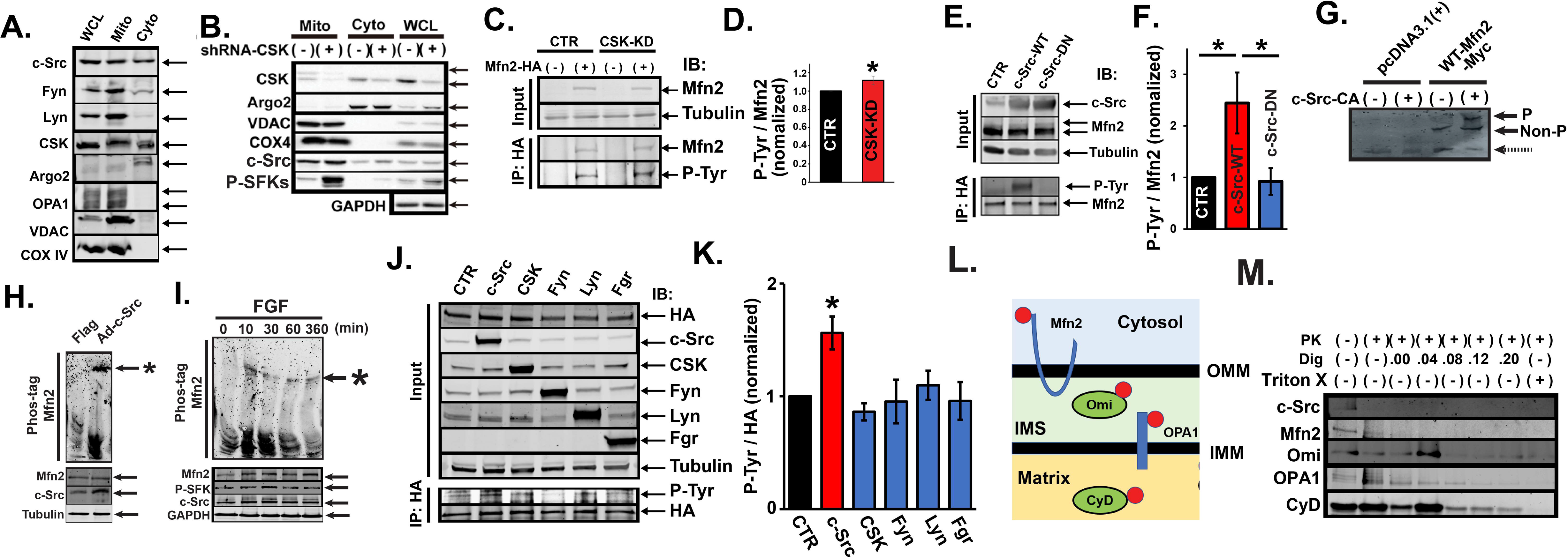
c-Src phosphorylates Mfn2. **A**. Expression of SFK members after mitochondrial fractionation of HEK293T cells. Argonaute 2 (Argo2) was used as a marker for the cytosolic fraction (Cyto). Voltage-dependent anion channel (VDAC), cytochrome c oxidase subunit 4 (COX4), and optic atrophy-1 (OPA1) were used as markers for the mitochondrial fraction (Mito). WCL, whole cell lysates. Representative images from 3 independent experiments. **B.** CSK-KD increases SFK activity in mitochondria. Representative immunoblots from HEK293T cells stably overexpressing CSK-shRNA (CSK-KD cells). Control cells (CTR) were prepared by overexpression of PLKO.1 empty vector. Immunoreactive bands were visualized by chemiluminescent immunoassay. Argo2 was used as a marker for the Cyto. VDAC and COX4 were used as markers for the Mito. Glyceraldehyde 3-phosphate dehydrogenase (GAPDH) was used for loading control for the WCL. **C.** Increased P-Tyr of Mfn2 in HEK293T cells by CSK knockdown. CTR and CSK-KD cells were transiently transfected with either Mfn2-HA or empty vector. Immunoprecipitation (IP) was performed with HA antibody, and P-Tyr of Mfn2 was detected by a general P-Tyr antibody. IB: immunoblot. **D.** Summary data of C (*n* = 4). Band intensity of immunoprecipitated P-Tyr was normalized to Mfn2 and expressed relative to CTR. \**P* < 0.05. **E.** Overexpression of c-Src increases P-Tyr of Mfn2. HEK293T cells stably overexpressing Mfn2-HA were transiently transfected with either empty vector (pcDNA3.1(+) as a control), c-Src-WT, or c-Src-DN. IP was performed with HA antibody, and P-Tyr of Mfn2 was detected by a general P-Tyr antibody. Band intensity of immunoprecipitated P-Tyr was normalized to Mfn2. **F.** Summary data of E (*n* = 4). \**P* < 0.05. **G.** P-Tyr of Mfn2 by c-Src-CA overexpression in HEK293T cells stably overexpressing WT-Mfn2-Myc assessed by 4% Phos-tag SDS-PAGE with Myc antibody. The phosphorylated form (P) and non-phosphorylated form (Non-P) are indicated by arrows. The bands indicated by dot arrows were likely non-specific bands since these also existed in the lysates from control cells. Representative images from 3 independent experiments. **H.** c-Src overexpression increases P-Tyr of endogenous Mfn2 in human cardiac fibroblasts assessed by 4% Phos-tag SDS-PAGE with Mfn2 antibody. Mfn2 band shifted by Phos-tag is shown with asterisk. Representative images from 3 independent experiments. **I.** FGF1 stimulation increases P-Tyr of endogenous Mfn2 in HCFs. **J.** Overexpression of c-Src, but not other SFK members, increases P-Tyr of Mfn2. HEK293T cells stably overexpressing Mfn2-HA were transiently transfected with either empty vector (pWZL-Neo-Myr-Flag-DEST as a control), c-Src, CSK, Fyn, Lyn, or Fgr. IP was performed with HA antibody, and P-Tyr of Mfn2 was detected by a general P-Tyr antibody. **K.** Summary data of I (*n* = 4). Band intensity of the immunoprecipitated P-Tyr was normalized by that in HA and expressed relative to CTR. \**P* < 0.05. **L.** Submitochondrial localizations of marker proteins for mitochondrial digestion assay. Red dots indicate the antibody binding sites. IMM, inner mitochondrial membrane; IMS, intermembrane space of mitochondria; Omi, human HTRA2; CyD cyclophilin D. **M.** Representative immunoblot patterns of mitochondrial digestion for submtiochodrial localization of c-Src. PK, Proteinase K; Dig, digitonin.

### c-Src increases ER-Mito interactions

Mfn2 has a key role in multiple mitochondrial and cellular functions (e.g., ER-Mito tethering, mitophagy, and ER stress) in addition to its canonical function as a mitochondrial fusion protein. The balance of these actions determines the mitochondrial shape, function, and distribution as well as the regulation of cell survival and death. Since we found that CSK-KD preferentially activates SFKs in the mitochondrial fraction (see **Fig. 1B**), we next explored the impact of P-Tyr of Mfn2 on these Mfn2 functions using CSK-KD cells.

First, ER-Mito interactions were assessed in live cells under confocal microscopy with a Förster resonance energy transfer (FRET)-based assay using OMM-targeted monomeric CFP (mt-CFP) and ER membrane-targeted monomeric YFP (ER-YFP) as a donor and an acceptor, respectively (**Fig. 2A**). Expression of either mt-CFP or ER-YFP alone does not show a significant FRET signal in HEK293T cells; their co-expression is required (**Fig. 2B**). In HEK293T cells with a stable knockdown of Mfn2 (Mfn2-KD), FRET between mt-CFP and ER-YFP was significantly lower than in control cells (**Fig. 2C-E**). Using this system, we found that FRET is significantly higher in CSK-KD cells compared to control cells (**Fig. 2F and G**), suggesting that the interaction between ER and mitochondria is enhanced in CSK-KD cells compared to control. We also biochemically confirmed that MAM increased in the CSK-KD cells compared to control by assessing the amount of ER membrane proteins IP_3_ receptor isoform 1 (IP_3_R1) in the mitochondrial fraction (**Appendix Fig. S2**). To further assess whether the increased ER-Mito interaction is mediated by SFK activity, we pre-incubated the cells with the SFK inhibitor PP2 (30 µM, 30 min). Pretreatment of PP2 for 30 min significantly decreased SFK activity in CSK-KD cells compared to control cells (**Appendix Fig. S1**). In addition, PP2 was able to abolish the observed increase in FRET in CSK-KD cells, indicating that these FRET changes are mediated via SFK activity (**Fig. 2H**). To further examine whether c-Src is involved in this mechanism, we genetically manipulated c-Src activity by overexpressing Src-WT and –DN (see also, **Fig. 1F**). We found that FRET increased after c-Src-WT but not c-Src-DN overexpression compared to control cells transfected with empty vector (**Fig. 2I**). These results indicate that increased FRET in CSK-KD cells is mainly through c-Src activation. Moreover, we showed that overexpression of Src-WT in Mfn2-KD does not increase FRET (**Fig. 2J**), suggesting that the action of c-Src on ER-Mito interactions is mediated via Mfn2. Lastly, we tested whether the c-Src-Mfn2 pathway observed in HEK293T cells also exists in primary cells. c-Src overexpression in HCFs (see also **Fig. 1H**) increased ER-Mito interactions similar to our observation in HEK293T cells, assessed by indirect immunofluorescence for FRET (Konig, Krasteva et al. 2006) (**Fig. 2K and L**).

**Fig. 2.**
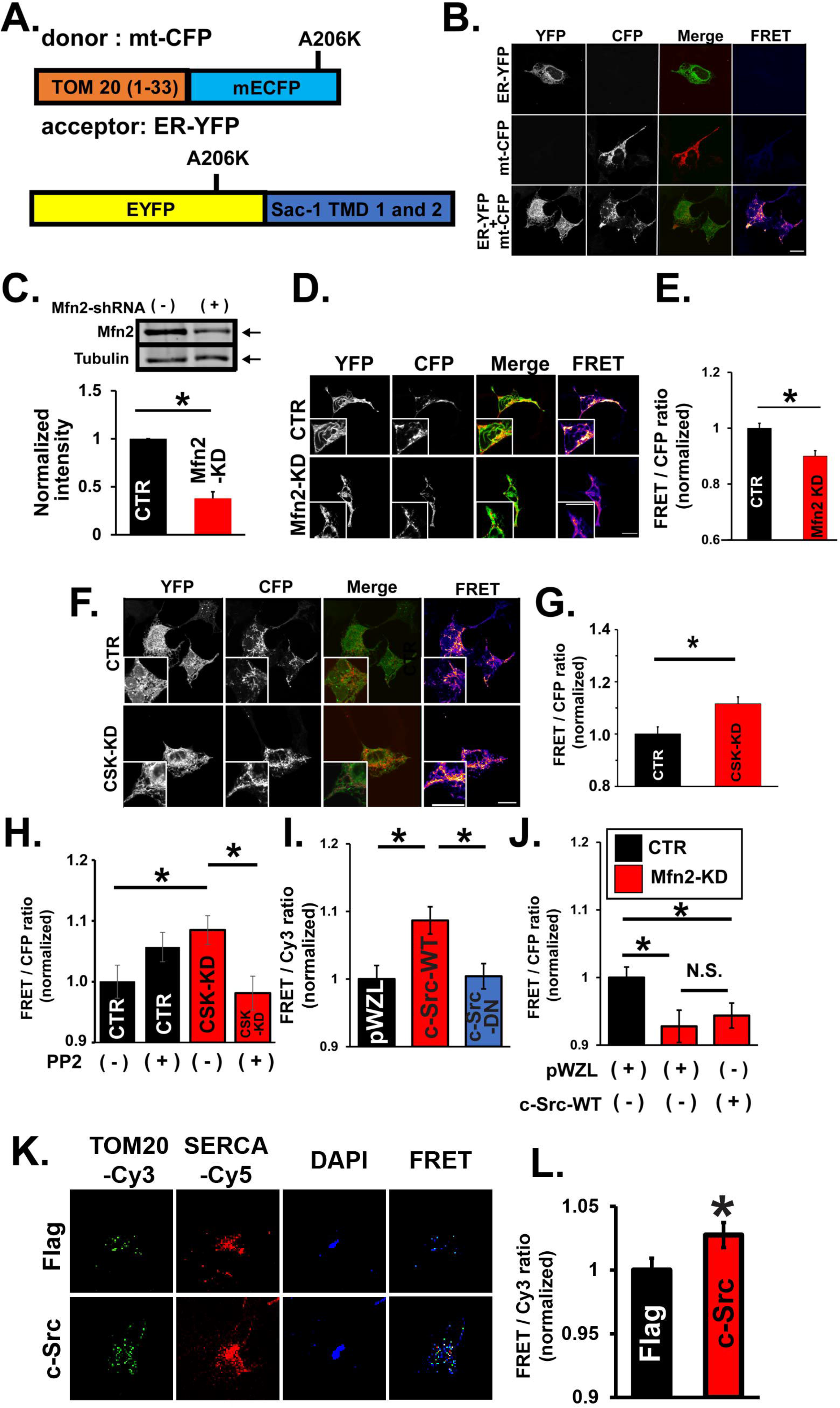
c-Src enhances ER-Mito interaction. **A**. Schematic diagram of a FRET donor and acceptor for assessing ER-mitochondria interaction. TMD, transmembrane domain. **B.** Expression of FRET sensors in HEK293T cells. Representative images of FRET detection in HEK293T cells transiently transfected with mt-CFP, ER-YFP, or both constructs. Scale bar, 20 µm. **C.** Establishment of an MFN2-KD cell line. *(Top):* Representative immunoblots of whole cell lysates obtained from HEK293T stably overexpressing Mfn2-shRNA (Mfn2-KD) or PLKO.1 (control; CTR). *(Bottom):* Summary data (*n* = 3). Mfn2 band intensity was normalized by that in tubulin and expressed relative to CTR. \**P* < 0.05. **D.** Mfn2 knockdown decreases ER-mitochondria interaction. Representative images of FRET detection in Mfn2-KD or CTR cells transfected with mt-CFP and ER-YFP. Scale bar, 20 µm. **E.** Summary of normalized FRET/CFP ratio from panel D (*n* = 104 and 100, for CTR and Mfn2-KD, respectively). The values were expressed relative to CTR. \**P* < 0.05. **F.** CSK-KD decreases ER-Mito interaction. Representative images of FRET detection in CSK-KD or CTR cells transfected with mt-CFP and ER-YFP. Scale bar, 20 µm. **G.** Summary of normalized FRET/CFP ratio from panel E (*n =* 132 and 135, for CTR and Mfn2-KD, respectively). The values were expressed relative to CTR. \**P* < 0.05. **H**. Effect of a SFK inhibitor PP2 on FRET/CFP ratio. CSK-KD or CTR HEK293T cells were co-transfected with FRET biosensors and treated with PP2 (30 µM) or DMSO (vehicle) for 30 min before FRET measurements (*n* = 58, 50, 54, and 26 for CTR+DMSO, CTR+PP2, CSK-KD+DMSO, and CSK-KD+PP2, respectively). The values were expressed relative to CTR+DMSO. \**P* < 0.05. **I.** Effect of a c-Src on FRET/CFP ratio. c-Src-WT, c-Src-DN, or pWZL empty vector (as a CTR) were co-transfected with FRET biosensors into HEK293T cells (*n* = 57, 97, and 82 for CTR, c-Src-WT and c-Src-DN respectively). The values were expressed relative to CTR. \**P* < 0.05. **J.** Effect of c-Src on FRET/CFP ratio in Mfn2-KD cells. c-Src-WT, or pWZL (as a control) were co-transfected with FRET biosensors into HEK293T cells. (*n =* 110, 117, and 118 for CTR, c-Src-WT and c-Src-DN respectively). \**P* < 0.05. **K.** Detection of close association between OMM protein TOM20 and ER membrane protein SERCA by indirect immunofluorescence and FRET in human cardiac fibroblasts (HCFs). Fluorescent images of donor (TOM20 labeled with Cy3-conjugated secondary antibody), acceptor (SERCA labeled with Cy5-conjugated secondary antibody) and FRET in HCFs infected with c-Src or Flag (as a control). **L.** Summary data of K (*n* = 20 and 26). The values were expressed relative to Flag. \**P* < 0.05.

To further investigate the ultrastructural alterations of ER-Mito contact sites by SFKs especially by c-Src, we assessed the ultrastructure of ER and mitochondria using CSK-KD cells by TEM (**Fig. 3A**). Importantly, the average distance between the ER membrane and OMM is significantly shorter in CSK-KD cells compared to control without a change in the average number of contact sites in each mitochondrion (i.e., number of associated ERs in each mitochondrion) or the morphology of mitochondrial cross-section (i.e., averaged aspect ratio [AR] of mitochondrial cross-section) (**Fig. 3B and C**). However, the ER–Mito interface length measured from each mitochondrial cross section (**Fig. 3A and B**) was not altered in CSK-KD cells compared to control cell (**Fig. 3C**). Combined with the results from live cell imaging, these results from TEM images suggest that SFK activation shortens the ER-Mito distance, possibly via c-Src-dependent Mfn2 phosphorylation, but does not significantly alter the interface length or the number of contact sites for each mitochondrion.

**Fig. 3.**
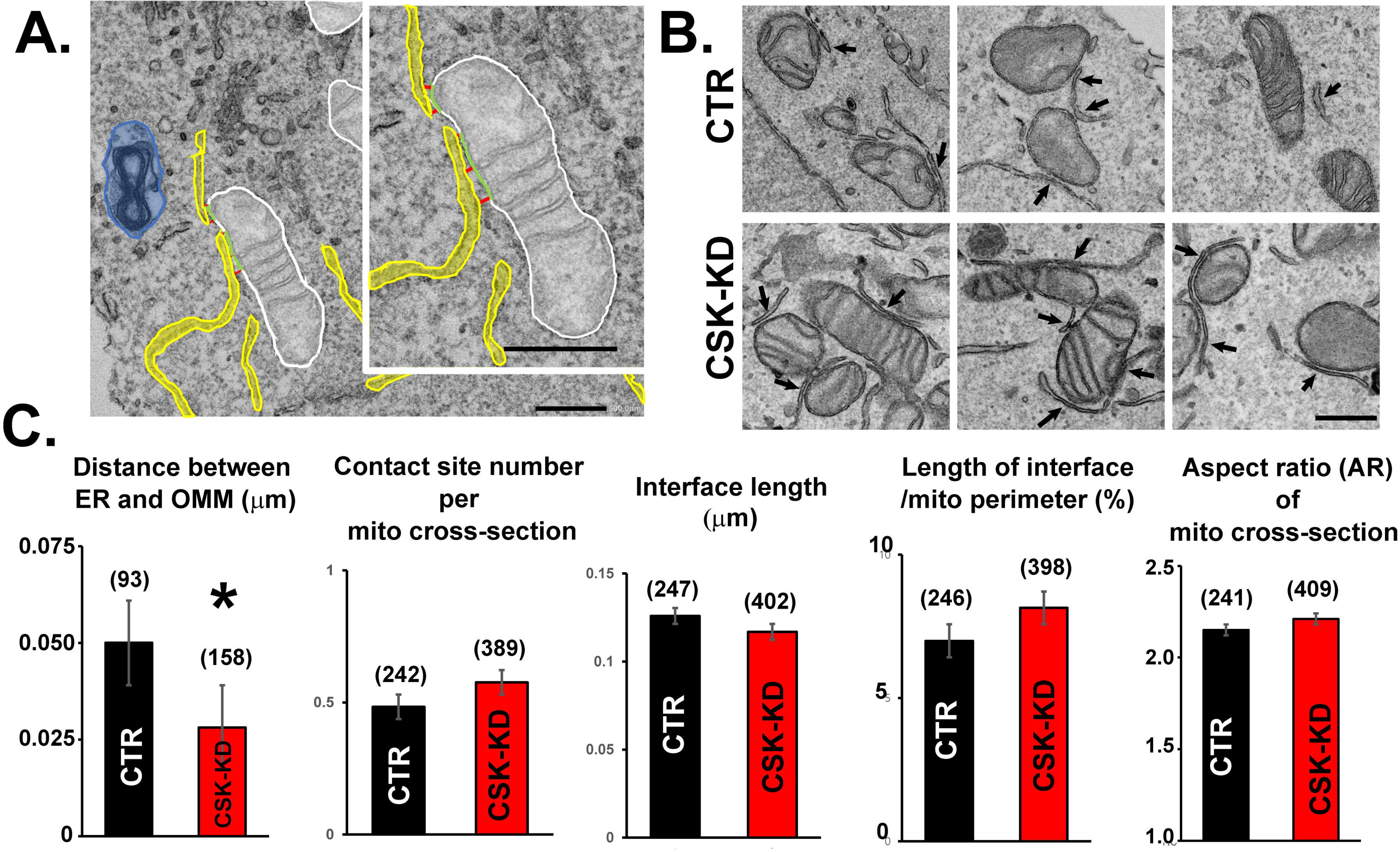
SFK activation shortens the distance between ER and mitochondria. **A**. Representative images of transmission electron microscopy (TEM) in HEK293T cells. White lines, green lines, yellow area, and blue area represent OMM, MAM interface, ER area, and mitophagosome area, respectively. MAM distances from three points (Red lines) were averaged to estimate the distance between ER and OMM. Scale bars, 500 nm. **B.** Representative TEM images in CSK-KD and CTR cells. Scale bar, 500 nm. Arrows point to the ER and mitochondria interaction sites. **C.** Summary data from B. Number in parentheses indicates the total number of images analyzed in each group \**P* < 0.05.

We further investigated whether SFK activity including c-Src influences other major Mfn2 functions, such as enhancing mitochondrial fusion, mitophagy, and ER stress. No significant difference was observed in mitochondrial shape and networks (**Fig. EV2A and B**), expression levels of major fission/fusion proteins (**Fig. EV1A and C**), autophagic flux (**Fig. EV2C**), and the numbers of mitophagosomes/autophagosomes (**Fig. EV2D**) in CSK-KD cells. Lastly, major ER stress markers, Grp94, Grp78, and CHOP (Ngoh, Papanicolaou et al. 2012) were assessed in CSK-KD and control cells (**Fig. EV2E**). We did not see any significant increase in Grp94, Grp78, and CHOP in CSK-KD cells. Together, these results indicate that SFK activation does not have a significant impact on other major functions of Mfn2.

### c-Src facilitates ER-Mito Ca^2+^ transport

We next investigated whether c-Src-mediated modification of the contact site influences ER-to-Mito Ca^2+^ transport. To measure the changes in Ca^2+^ concentration at the mitochondrial matrix ([Ca^2+^]_mt_) in response to cytosolic Ca^2+^ elevation, we used the mitochondrial matrix-targeted Ca^2+^-sensitive biosensor mtRCamp1h (Hamilton, Terentyeva et al. 2018) (**Fig. 4A**). c-Src-overexpression cells started to increase [Ca^2+^]_mt_ immediately after IP_3_R-mediated ER Ca^2+^ release by ATP treatment, which stimulates endogenous G_αq/11_ protein-coupled P2Y receptor (Schrage, Schmitz et al. 2015). Average time-to-peak-[Ca^2+^]_mt_ became significantly shorter and the magnitude of peak [Ca^2+^]_mt_ significantly increased in c-Src-overexpression cells compared to control cells (**Fig. 4A and B**) without affecting the size or rate of cytosolic Ca^2+^ concentration ([Ca^2+^]_cyto_) elevation (**Fig. 4C and D**). CSK-KD cells showed the similar mtCa^2+^ uptake profile by ATP treatment (**Fig. EV3**).

**Fig. 4.**
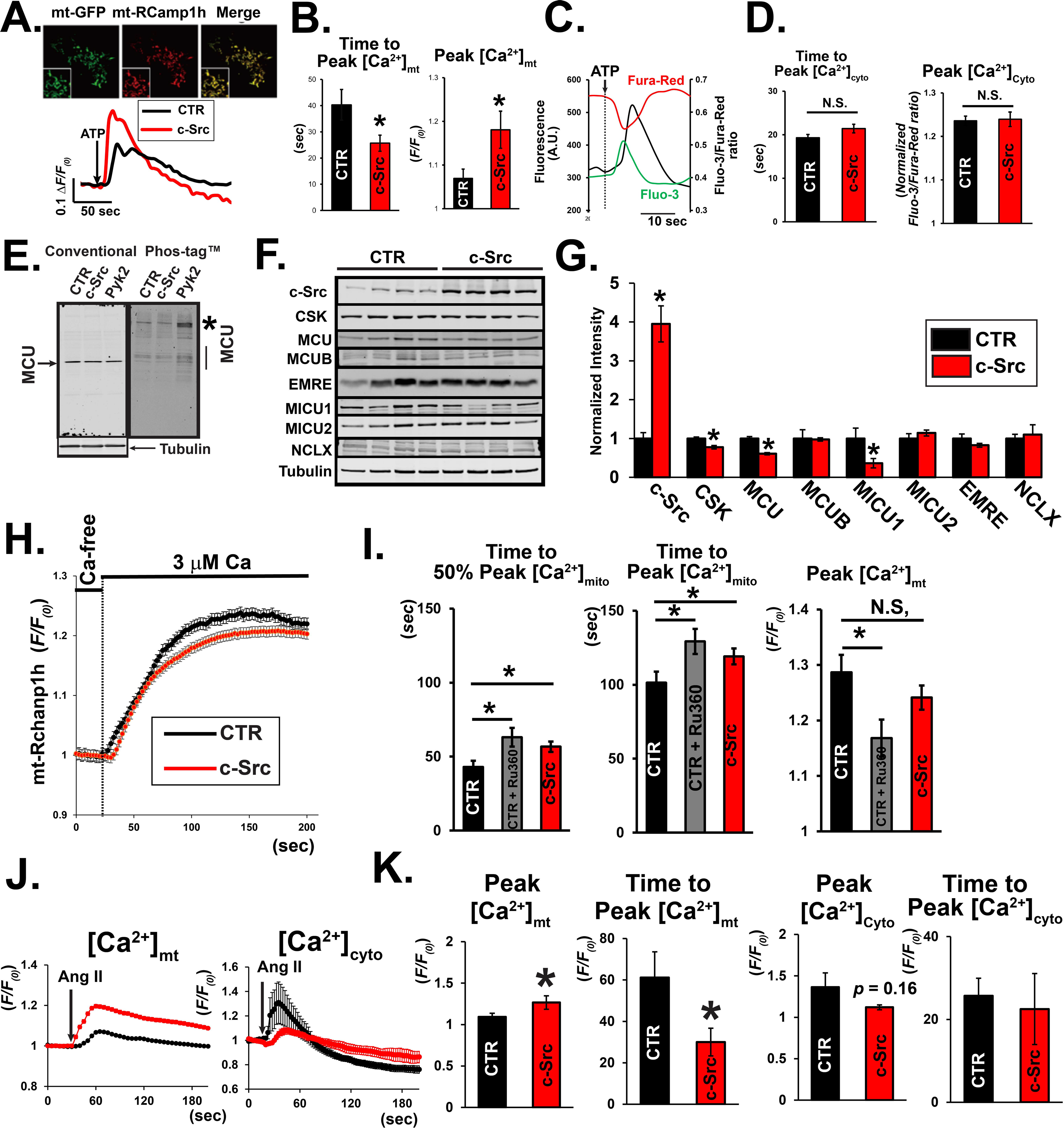
c-Src increases mtCa^2+^ uptake in response to cytosolic Ca^2+^ elevation. **A. *Top:*** Expression of mitochondrial matrix-targeted Ca^2+^-sensitive biosensor, mt-RCaMP1h, in HEK293T cells co-transfected with mitochondrial matrix-targeted GFP (mt-GFP). ***Bottom:*** Representative traces of changes in Ca^2+^ concentration in the mitochondrial matrix ([Ca^2+^]_mt_) in HEK293T cells stably overexpressing c-Src and pcDNA3.1(+) empty vector (CTR) in response to 2 mM ATP. [Ca^2+^]_mt_ was assessed by mt-RCamp1h. **B.** Summary data of A (*n =* 20 and 35 for CTR and c-Src overexpressing cells, respectively). \**P* < 0.05. **C.** Representative traces of changes in cytosolic Ca^2+^ concentration ([Ca^2+^]_cyto_) in single HEK293T cells in response to 2 mM ATP. Changes in [Ca^2+^]_cyto_ were measured from a live single cell using a ratiometric method (black line) with two Ca^2+^-sensitive dyes, Fluo-3 (green line) and Fura-Red (red line), under confocal microscopy. **D.** Summary data of cytosolic Ca^2+^ handling profile in CTR and c-Src overexpressing cells (*n* = 209 and 202 for CTR and c-Src cells, respectively). N.S., not significant. **E.** P-Tyr levels of MCU after c-Src and Pyk2 overexpression in HEK293T cells stably overexpressed MCU-flag assessed by Phos-tag SDS-PAGE. Conventional SDS-PAGE is shown for comparison. The location of the phosphorylated MCU shifted by Phos-tag is shown with an asterisk. **F.** Protein expression profile in CTR and c-Src overexpressing cells. **G.** Summary data of F. Each band intensity was normalized by tubulin. *n* = 4 for each group. \**P* < 0.05. **H.** Averaged traces of the changes in [Ca^2+^]_mt_ in response to treatment with 3 μM Ca^2+^ in permeabilized CTR and c-Src-overexpressing HEK293T cells transfected with mt-RCamp1h. *N* = 51 and 84 for CTR (black) and c-Src cells (red), respectively. **I.** Summary data of H. The data from CTR cells treated with MCU blocker 1 μM Ru360 (gray bars, *n* = 27) was shown as a comparison. \**P* < 0.05. **J.** Effect of c-Src overexpression on mitochondrial and cytosolic Ca^2+^ handling in response to 10 μM angiotensin II (Ang II) stimulation in human cardiac fibroblasts (HCFs) assessed by mtRhamp1h and Fluo-3, respectively. (*n =* 19 and 17 for [Ca^2+^]_mt_, and *n =* 15 and 14 for [Ca^2+^]_cyto_). **K.** Summary data of [Ca^2+^]_mt_ and [Ca^2+^]_cyto_ from panel J. \**P* < 0.05.

We and others previously showed the phosphorylation of MCU, a pore-forming subunit of mtCUC, by protein kinases (O-Uchi, Jhun et al. 2014, Lee, Min et al. 2015, Xie, Song et al. 2018, Zhao, Li et al. 2019) including another non-receptor-type PTK Pyk2 (O-Uchi, Jhun et al. 2014). However, c-Src was not able to phosphorylate MCU, as assessed by the Phos-tag SDS-PAGE (Kinoshita, Kinoshita-Kikuta et al. 2006) (**Fig. 4E**, see also **Fig. 1G-I**), indicating that the c-Src-dependent mtCa^2+^ uptake upregulation is likely not via MCU phosphorylation. The expression levels of mtCa^2+^ influx/efflux-related proteins (i.e., mtCUC subunit proteins and mitochondrial Na^+^/Ca^2+^ exchanger [NCLX]) between control and c-Src overexpression cells were assessed. A significant decrease in the pore-forming subunit MCU and one of the regulatory subunits MICU1 was detected in c-Src overexpression cells, despite of mtCa^2+^ facilitation in these cells (**Fig. 4F and G**). The MICU1/MCU ratio did not significantly alter between two groups (1.026 ± 0.28 and 0.612 ± 0.20, in CTR and c-Src overexpression cells, respectively, *p*=0.27). CSK-KD cells also showed a slight (but not significant) decrease in MCU and a significant decrease in MICU1 (**Fig. EV1**).

To further investigate the detailed mechanism of how c-Src alters the mtCa^2+^ uptake profile, we assessed the activity of mtCUC in permeabilized cells where the ER-Mito interactions have been disrupted. After digitonin permeabilization, the mtCa^2+^ uptake was initiated by switching the medium from a buffer mimicking the cytosolic ionic composition (termed “intracellular buffer”) with EGTA (Ca^2+^-free) to the intracellular buffer with 3 μM free Ca^2+^ (Raffaello, De Stefani et al. 2013, De Stefani, Raffaello et al. 2011) (**Fig. 4H**). Treatment with a MCU blocker (Ru360, 1 μM) significantly slowed down the speed of mtCa^2+^ uptake and decreased mtCa^2+^ plateau level in the control cells, confirming that the majority of mtCa^2+^ uptake is via mtCUC (**Fig. 4I**). The c-Src-overexpressing cells showed significantly slower mtCa^2+^ uptake, indicating that mtCUC activity in c-Src-overexpressing cells is lower than control cells (**Fig. 4H and I**), which is consistent with our observations in the mtCUC protein expression profile (i.e., decreased in a pore-forming protein MCU) (**Fig. 4G&H**). CTR and c-Src overexpressing cells reached a similar plateau level as control cells (**Fig. 4H&I**).

Lastly, we tested whether the c-Src-Mfn2 pathway observed in HEK293T cells also affects ER-Mito Ca^2+^ transport in primary cells. We measured mtCa^2+^ uptake in response to angiotensin II (Ang II) stimulation, which stimulates endogenous G_αq/11_ protein-coupled AT1 receptor in HCFs and induces ER Ca^2+^ release (Mohis, Edwards et al. 2018). Average time-to-peak-[Ca^2+^]_mt_ became significantly shorter and the magnitude of peak [Ca^2+^]_mt_ significantly increased in c-Src-overexpression HCFs compared to control HCFs (**Fig. 4J and K**), which is similar to our results in HEK293T cells. The greater mitochondrial response was not secondary to alterations of the cytosolic response because we observed a trend towards a reduction in the [Ca^2+^]_cyto_ transient that was not significant (**Fig. 4J and K**).

Taken together, we demonstrate that c-Src activates the mtCa^2+^ uptake profile (increased peak [Ca^2+^]_mt_ and faster mtCa^2+^ uptake) without affecting cytosolic Ca^2+^ handling, even though mtCUC channel activity is reduced, possibly because of a decrease in the number of mtCUC channels. The activation of mtCa^2+^ uptake observed in the intact cells after c-Src activation is unlikely via mtCUC channel activation by c-Src (i.e., c-Src-dependent P-tyr of MCU). We conclude instead that mtCa^2+^ uptake is facilitated by the increased efficiency of ER-to-Mito Ca^2+^ transport by decreasing the ER-Mito distance by c-Src.

### c-Src phosphorylates the C-terminal tail of Mfn2

We next attempted to identify the c-Src-specific phosphorylation site(s) in Mfn2 and investigated their roles in ER-Mito tethering. Human Mfn2 contains 15 tyrosine residues: ten in the N-terminus, two in the C-terminus, and three tyrosine residues located in the transmembrane domains. We found that four of these residues (i.e., Y49, Y61, Y269, and Y752) in human Mfn2 are predicted to be phosphorylated by known PTKs and/or c-Src based on two phosphorylation prediction programs, NetPhos 3.1 (Blom, Gammeltoft et al. 1999), and GPS 5.0 (Wang, Xu et al. 2020) (**Fig. 5A**). In mass spectrometry, Y448 was the only site reported to be phosphorylated in mouse Mfn2 (Hornbeck, Kornhauser et al. 2012). Importantly, all five of these sites reside in the cytoplasmic face of the Mfn2 structure, based on the conventional Mfn2 topological model (Koshiba, Detmer et al. 2004, Chen, H., Detmer, Ewald, Griffin, Fraser, and Chan 2003) (**Fig. 5A**). Thus, we selected these five Tyr residues (Y49, Y61, Y269, Y448, and Y752) as candidates for c-Src-specific P-Tyr sites. Among these sites, Y61 does not exist in rodent Mfn2 (**Appendix Fig. S3A**). In contrast, the sequences surrounding Y49, Y269, Y448, and Y752 are well-conserved across humans, rats, and mice (**Appendix Fig. S3A**). The homologous protein human Mfn1 (Franco, Kitsis et al. 2016) has Tyr residues corresponding to Mfn2-Y61 and –Y269, but not Y49, Y448, and Y752 (**Appendix Fig. S3B**).

**Fig. 5.**
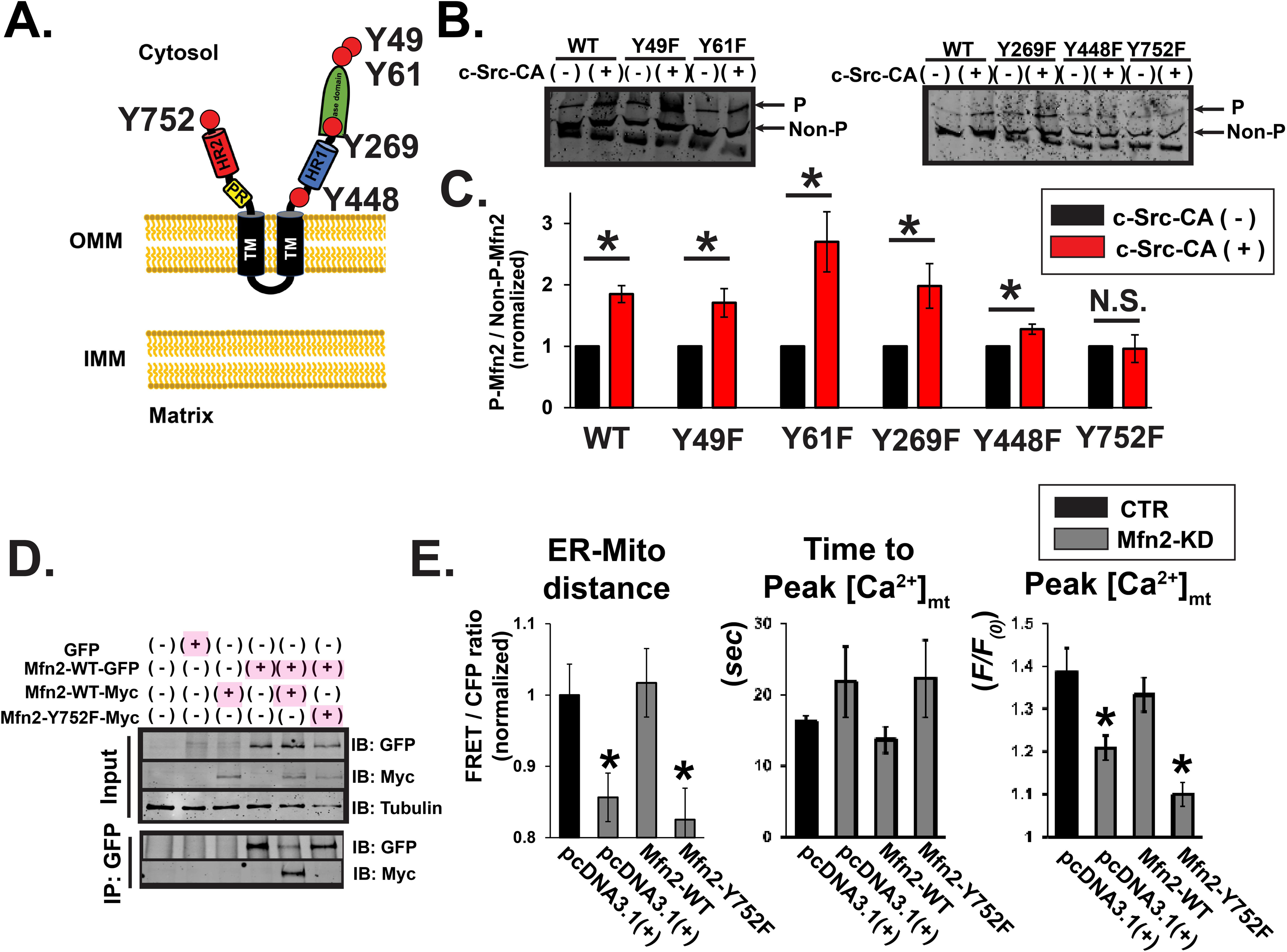
c-Src phosphorylates C-terminal tail of Mfn2. **A**. Candidate of c-Src-specific sites in human Mfn2. **B.** Phosphorylated Mfn2 by c-Src-CA in HEK293T cells stably overexpressing WT or mutant-Mfn2 assessed by 4% Phos-tag SDS-PAGE with Myc antibody (see also **Fig. 1G**). The phosphorylated form (P) and non-phosphorylated form (Non-P) are indicated by arrows. **C.** Summary data of P-Mfn2/ Non-P-Mfn2 ratio calculated from B. Phospho (P) / non-phospho (Non-P) ratio in each Mfn2 mutant after c-Src-CA overexpression was normalized to that without c-Src-CA overexpression. \**P* <0.05. N.S., not significant. **D.** Co-IP of GFP– and myc-tagged Mfn2 expressed in Mfn2-KD cells. **E. *Left:*** Reintroduction of WT-Mfn2, but not –Y752F, into Mfn2-KD cells decreases ER-Mito interaction (*n* 53, 86, 66, 82). \**p*<0.05 compared to control (CTR, black). ***Middle and Right:*** Reintroduction of WT-Mfn2, but not –Y752F, into Mfn2-KD cells increases the speed and magnitude of mtCa^2+^ uptake (*n =* 17, 39, 19, 11).

We next created HEK293T cells stably overexpressing Myc-tagged dephospho-mimetic mutants of Mfn2 (i.e., mutated Tyr [Y] residue to phenylalanine [F]) and transfected them with c-Src-CA. P-Tyr of Mfn2 by c-Src-CA was assessed by Phos-tag SDS-PAGE (see also **Fig. 1G**). In cells expressing Mfn2-WT-, Mfn2-Y49F, –Y61F, – Y269F, and –Y448F, c-Src increased P-Tyr of Mfn2 (i.e., increased the ratio of phosphorylated and non-phosphorylated Mfn2 bands) (**Fig. 5B and C**), suggesting that these sites are likely not c-Src-specific sites. Increased P-Tyr levels in Mfn2-Y448F were smaller than those in other mutants and WT. In contrast, cells expressing Mfn2-Y752F did not show a significant increase in P-Tyr of Mfn2 by c-Src-CA, indicating that Y752 is a potential c-Src-specific P-Tyr site in Mfn2 (**Fig. 5B and C**). Moreover, Mfn2-Y752F failed to associate with WT-Mfn2 as assessed by Co-IP (**Fig. 5D**), indicating that P-Tyr of Y752 is required for Mfn2-Mfn2 dimerization.

To further investigate the impact of the Y752F mutation on the microdomain size and mtCa^2+^ uptake profile, we reintroduced WT or Mfn2-Y752F into Mfn2-KD cells (see also **Fig. 2C**). Reintroduction of Mfn2-WT successfully normalized the distance between ER and mitochondria, and the mtCa^2+^ uptake profile in Mfn2-KD cells was similar to levels comparable to the control cells, but Mfn2-Y752F failed to show these changes in Mfn2-KD cells (**Fig. 5E**). The data suggests that Y752 at the C-terminal tail of Mfn2 is a potential c-Src-specific site that is required for Mfn2-Mfn2 interaction and regulation of ER-Mito tethering.

### SFK activation modulates mitochondrial functions

We next investigated whether the increased efficiency of ER-Mito Ca^2+^ transport by c-Src dependent P-Tyr of Mfn2 impacts the downstream mitochondrial signaling. Excessive and chronic mitochondrial Ca^2+^ uptake increases mROS production from mitochondria possibly via changing mitochondrial membrane potentials (*Δψ_m_*) (Jhun, Mishra et al. 2016) and/or inhibiting complex I activity (Malyala, Zhang et al. 2019). We found increased c-Src activity by CSK-KD (see **Figs. EV1 and S1**) significantly depolarized *Δψ_m_* compared to control, assessed by *Δψ_m_*-sensitive dye tetramethylrhodamine ethyl ester (TMRE) (**Fig. 6A and B**), which can also decrease the driving force of Ca^2+^ entry into the mitochondria via mtCUC channel in addition to decreased MCU protein. We next measured mROS levels in CSK-KD cells using a mitochondrial matrix-targeted H_2_O_2_-sensitive biosensor mt-roGFP2-Orp1 (Meyer, Dick 2010). This pH-insensitive biosensor contains Orp-1, a peroxidase that catalyzes the oxidation of H_2_O_2_. When mt-roGFP2 and Orp-1 are coupled together, it creates an mROS-sensitive fluorescent biosensor. Increased mROS results in a decrease in mt-roGFP2-Orp1 fluorescence. Basal mROS levels and maximum mROS production were measured by stimulating the cells with an oxidative stress inducer, tert-butyl hydroperoxide (T-BH) followed by the application of a reducing agent, dithiothreitol (DTT) (**Fig. 6C**). CSK-KD cells had higher basal mROS levels as well as higher maximum mROS production compared to control cells (**Fig. 6D**). Consistent with increased mROS and depolarized *Δψ_m_*, we observed reduced basal and maximal respiration in CSK-KD cells compared to control (**Fig. 6E and F**). These results indicate that SFK activation modulates mROS and mitochondrial bioenergetics, possibly via c-Src-dependent Mfn2 phosphorylation.

**Fig. 6.**
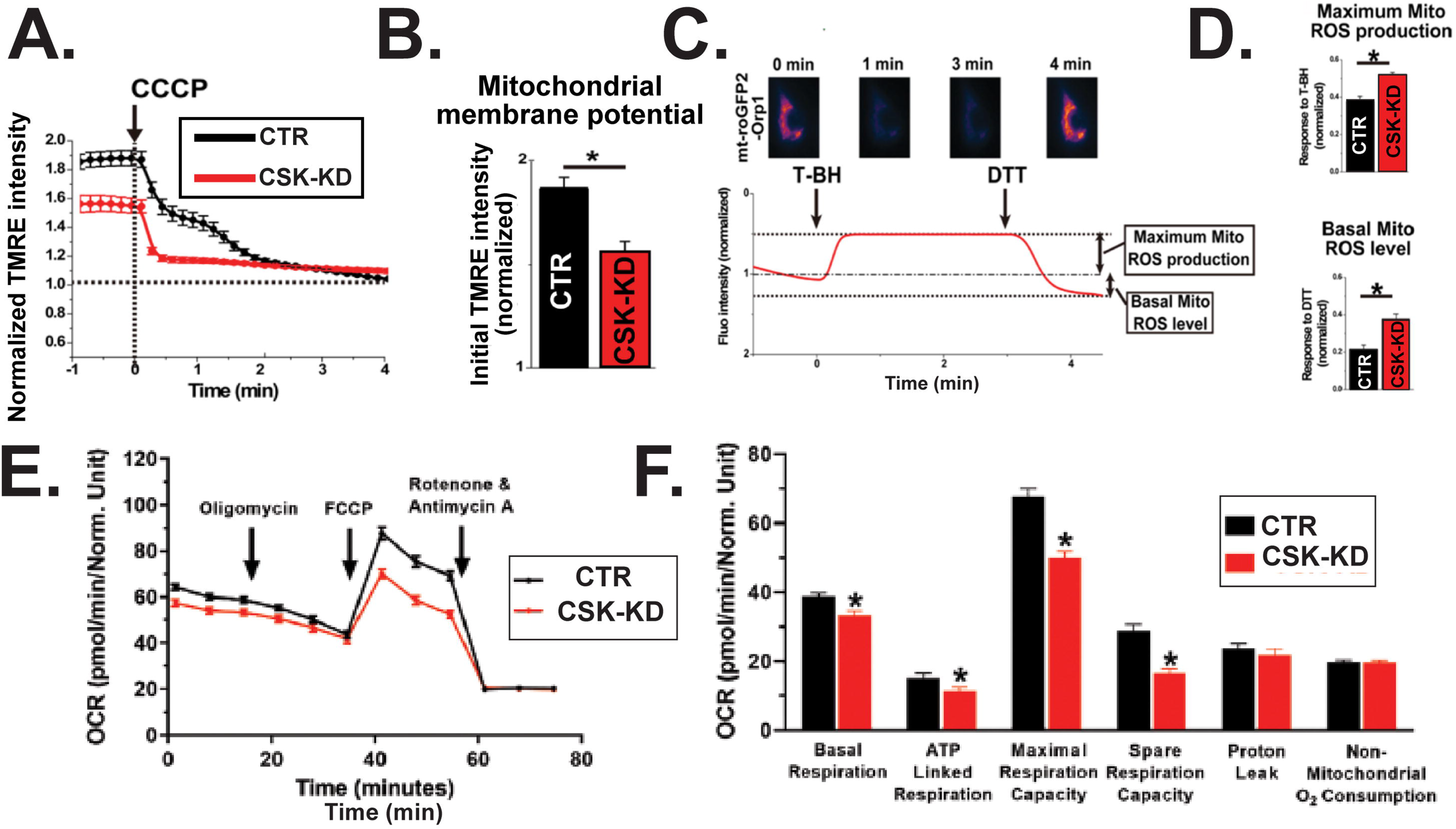
SFK activation modulates mitochondrial functions. **A**. Averaged traces of the changes in fluorescence intensity of tetramethylrhodamine, ethyl ester (TMRE) in response to treatment with CCCP in CTR and CSK-KD HEK293T cells. Initial TMRE intensity before CCCP treatment was normalized with the value after CCCP treatment. **B.** Summary data of A (*n* = 50 and 65 for CTR and CSK-KD cells, respectively). \**P* <0.05**. C.** Representative time-lapse images (top) and traces (bottom) of HEK293T cells transfected with mitochondrial targeted ro-GFP2-Orp1 (mt-roGFP2-Orp1) upon stimulation with an oxidative stress inducer, tert-butyl hydroperoxide (T-BH, 10 mM) and a reducing agent, dithiothreitol (DTT, 100 mM). **D.** Summary data of C (*n* = 32, and 22 for CTR and CSK-KD cells, respectively). \**P* <0.05. **E.** Oxygen consumption rate (OCR) measurements obtained over time (min) using an extracellular flux analyzer in CTR and CSK-KD cells. Data from 3 independent experiments. **F.** Summary data of E (*n* = 3 for each group). \**P* <0.05.

## Discussion

In this study, we identified a novel mechanism that regulates ER-Mito tethering, ER-Mito Ca^2+^ transport, and mROS production via P-Tyr of Mfn2 (**Fig. 7**). Our major findings are **1)** Mfn2 is a novel substrate of c-Src, one of the SFKs, in mitochondria; **2)** c-Src can phosphorylate the C-terminal tail of Mfn2, a domain critical for Mfn2 dimerization (Koshiba, Detmer et al. 2004, Chen, H., Detmer, Ewald, Griffin, Fraser, and Chan 2003) and redox sensing (Mattie, Riemer et al. 2018); and **3)** c-Src-dependent P-Tyr of Mfn2 at Y752 shortens the ER-Mito distance and facilitates ER-to-Mito Ca^2+^ transfer, followed by an increase in mROS generation. Since the homologous protein Mfn1 does not possess Y752, this tethering regulation by c-Src is likely Mfn2-specific. These results provide new insights into the molecular basis of the modulation of ER-Mito tethering machinery as well as mtCa^2+^ handling at MAM. Furthermore, this study unveils a new pathway that causes mtCa^2+^ overload and subsequent mROS elevation that could be targeted under pathological conditions.

**Fig. 7.**
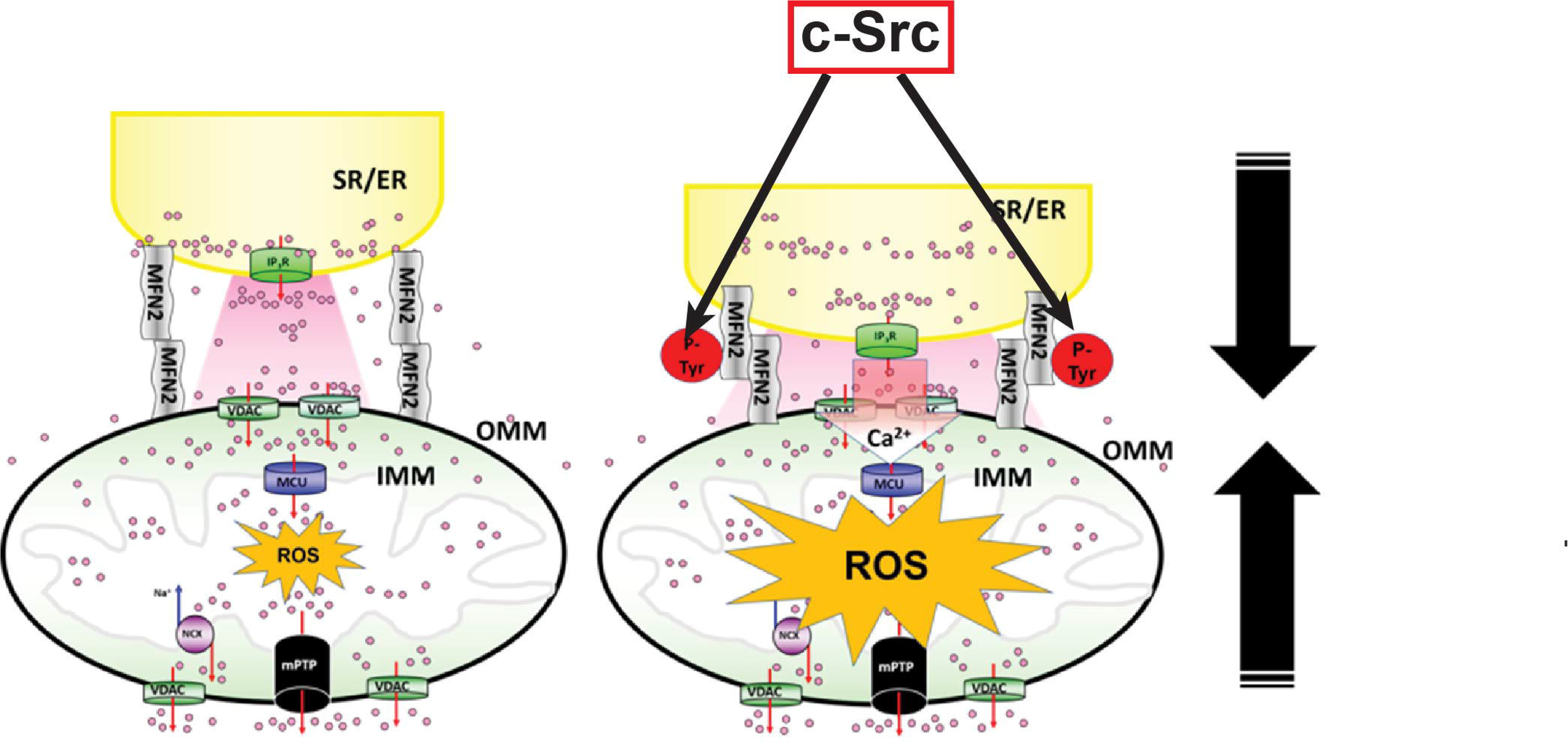
Proposed model: Src-dependent Mfn2 phosphorylation decreases the ER-Mito distance and facilitates mtCa^2+^ uptake and ROS generation.

### Molecular mechanism underlying the regulation of ER-Mito tethering by tyrosine phosphorylation of Mfn2

In this study, we show that c-Src-dependent Mfn2 phosphorylation at Y752 at its c-terminal tail can modulate ER-Mito tethering function (**Fig. 5**). Mitochondrial fission/fusion proteins including Mfn2 receive PTMs by kinases that are activated by cellular stress (Chang, Blackstone 2010), which may change the functional properties of these proteins including their sub-cellular/mitochondrial distribution, protein-protein interactions and the formation of higher-order assembly (e.g., dimerization in Mfn1/2), and their GTPase activity (Chang, Blackstone 2010, Adaniya, O-Uchi et al. 2019). Our data suggest that **1)** P-Tyr of Mfn2 does not increase the number of tethering sites, but likely changes the structure of existing tethering units (**Fig. 3**); and **2)** C-terminal Y752, which is located far from the GTPase domain, is the c-Src site that regulates Mfn2-Mfn2 dimerization and the tethering function (**Fig. 5**). This result is consistent with previous observations that the C-terminal heptad-repeat (HR) domain in Mfn1 and Mfn2 is critical for forming homotypic (Mfn1-Mfn1 or Mfn2-Mfn2) or heterotypic (Mfn1-Mfn2) dimers independent from their GTPase activity (Koshiba, Detmer et al. 2004, Chen, H., Detmer, Ewald, Griffin, Fraser, and Chan 2003), despite the two potential models of Mfn topology (i.e., the C-terminus facing either the cytoplasm or the intermembrane space of mitochondria [IMS]) (Mattie, Riemer et al. 2018).

The GTPase activity of Mfn2 is critical for fusing the mitochondria as well as generating Mfn2-Mfn2 tethering units, as evidenced by functional assessments of Mfn2 knockout cells (de Brito, Scorrano 2008, Naon, Zaninello et al. 2016) and in Mfn2 mutations found in Charcot-Marie-Tooth disease type 2A (CMT2A) (Zuchner, Mersiyanova et al. 2004). Increased Mfn2 GTPase activity is expected to increase mitochondrial fusion, generate new tethering units, as well as to shorten the distance between two organelles. This idea is also supported by the previous study from Csordas et al. that expressed synthetic ER-Mito linking molecules (drug-inducible fluorescent interorganelle linkers) in cultured cells (Csordas, Varnai et al. 2010). Since Y752 is located at the C-terminus of Mfn2 and is far from the GTPase domain, P-Tyr of Mfn2 at Y752 is unlikely to alter GTPase activity and therefore will not influence the number of tethering units. Instead, P-Tyr of Mfn2 at Y752 is more likely to change the length of existing tethering machinery, possibly through conformation changes of its 3D structure. This idea is also supported by the recent report that ubiquitination of Mfn2 by the E3 ubiquitin ligase parkin at K416, which is also located outside of GTPase domain, is required for maintaining the OMM-ER distance without changing Mfn2 GTPase activity (Basso, Marchesan et al. 2018).

Although Y448 is not predicted to be a c-Src site in Mfn2, we found that the c-Src-induced increase in P-Tyr levels of Mfn2 was smaller in Mfn2-Y448F-expressing cells compared to Mfn2-WT-expressing cells (**Fig. 5**). Since P-Tyr at Y448 is detectable by mass spectroscopy from mouse samples (Hornbeck, Kornhauser et al. 2012), basal P-Tyr levels in Y448 may affect the level of P-Tyr of Mfn2 by c-Src, possibly serving as a priming site for causing c-Src-dependent P-Tyr of Mfn2. Other tyrosine kinases at the OMM (e.g., the two SFK members we detected in the mitochondria, see **Fig. 1A**) and/or mitochondria-localized tyrosine phosphatases (Miyazaki, Tanaka et al. 2006, Miyazaki, Neff et al. 2003, Tibaldi, Brunati et al. 2008) may also regulate the basal P-Tyr of Y488 in Mfn2.

In summary, our data show that Mfn2 is a novel c-Src substrate in the mitochondria and c-Src-dependent Mfn2 phosphorylation at the C-terminal tail decreases the ER-Mito distance, possibly by modulating the 3D structure of the existing ER-Mito tethering units, rather than increasing the number of tethers.

### Molecular mechanism underlying the regulation of ER-Mito Ca^2+^ transport by tyrosine phosphorylation of Mfn2

Our data suggest that c-Src-dependent P-Tyr of Mfn2 increases the efficiency of ER-to-Mito Ca^2+^ transport without significantly affecting cytosolic Ca^2+^ handling, mainly by changing the distance between ER and mitochondria despite reduced mtCUC activity (**Figs. 4-6**). Our TEM data showed that c-Src shortens the ER-Mito distance from ∼50 nm down to ∼25 nm without changing the perimeter of MAM (**Fig. 3**). However, these effects are much smaller than those reported in cell models expressing synthetic ER-OMM linker molecules (Csordas, Renken et al. 2006, Csordas, Varnai et al. 2010). Csordas et al. showed that artificial ER-Mito tethering increases the perimeter of MAM (up to ∼1 μm), and decreases the distance between the ER and OMM (down to ∼6 nm), followed by the enhancement of Ca^2+^ transport from ER to mitochondria (Csordas, Renken et al. 2006, Csordas, Varnai et al. 2010).

At the resting state, the electrochemical driving force for mtCa^2+^ uptake via the mtCUC is provided by *Δψ_m_* across the IMM. The mtCa^2+^ uptake rate is very slow near the resting [Ca^2+^]_c_, but rapidly increases if the [Ca^2+^]_c_ in the proximity of the mitochondria reaches around 10-20 μM (Csordas, Thomas et al. 1999, Csordas, Varnai et al. 2010, Rizzuto, Pinton et al. 1998, Giacomello, Drago et al. 2010). When IP_3_R releases ER Ca^2+^ to the MAM, the Ca^2+^ concentration at the microdomain ([Ca^2+^]_MAM_) exceeds 10 μM (Csordas, Varnai et al. 2010), which is sufficient to trigger mtCa^2+^ uptake via MCU. After a decrease in tethering length, the MAM region becomes smaller, which results in shorter time-to-peak-[Ca^2+^]_MAM_ and larger peak [Ca^2+^]_MAM_ (Csordas, Varnai et al. 2010). A recently published mathematical model from Qi et al. suggests that the concentration of Ca^2+^ that is sensed by MCU in MAMs depends on the distance between MCU and the point source (i.e., IP_3_Rs at the ER membrane) and the rate of Ca^2+^ diffusion across the MAM (Qi, Li et al. 2015). This mathematical model provides several key insights that help explain our experimental results. They they proposed that ∼30 nm distance between IP_3_Rs and mtCUC is the critical range for mitochondria to efficiently uptake and modulate Ca^2+^ from MAM and to modulate cytosolic Ca^2+^ signaling. When the distance is ∼ 30 nm, about 50% of the ER-released Ca^2+^ ions are taken up by mitochondria. If the distance becomes < 30 nm, a sharp increase in [Ca^2+^]_MAM_ is predicted, and the majority of Ca^2+^ ions released by the ER are captured by the mitochondria and this effect reaches the maximum around 20 nm. However, at a very short distance (< 20 nm, e.g., in the case of artificial ER-OMM linker molecules (Csordas, Renken et al. 2006), the ER-released Ca^2+^ ions almost saturate both mitochondria and the cytosol, thus the total amount of ER-released Ca^2+^ decreases. Given that OMM and IMM distance is ∼8 nm (Herrmann, Neupert 2004), the estimated distance between IP_3_Rs and mtCUC after CSK-KD is around 30-35 nm, which strongly supports the physiological relevance of P-Tyr of Mfn2-mediated regulation of ER-Mito distance under c-Src activation. The magnitude of change induced by c-Src overexpression or knocking down of the SFK inhibitor CSK is sufficient for maximizing the efficiency of ER-Mito Ca^2+^ transport that can change [Ca^2+^]_MAM_, and subsequently [Ca^2+^]_m_ despite having lower mtCUC channel activity compared to control cells (**Fig. 4 and EV3**). While cells expressing artificial ER-Mito tethers can produce narrower macrodomains (∼ 6 nm) between ER and mitochondria (Csordas, Renken et al. 2006, Csordas, Varnai et al. 2010), the increase in ER-Mito Ca^2+^ transport induced by artificial ER-Mito tethers possibly derives from the increased the length of the MAM perimeter rather than the ER-Mito distance.

In summary, our data show that P-Tyr of Mfn2 by c-Src is capable of enhancing the efficiency of ER-Mito Ca^2+^ transport by shifting the ER-Mito distance to the “sweet spot” for gaining the efficiency of ER-Mito Ca^2+^ transport. Reduction in mtCUC activity via decreased expression of subunit proteins and depolarized *Δψ_m_* might be a compensatory mechanism for protecting the mitochondrial from mtCa^2+^ overload. MCU expression may be partly regulated transcriptionally, but the changes in MICU1 are likely via a post-translational mechanism (**Fig. EV4**). Further studies are required for understanding the detailed mechanism of how c-Src signaling decrease mtCUC protein expressions.

### Role of c-Src signaling in regulating mitochondrial functions

The experiments in this study have revealed that c-Src not only modulates the MAM space size and the profile of mtCa^2+^ transport (**Figs. 2-5 and 7**), but also changes the profile of *ΔΨ_m_*, mROS, and mitochondrial respiration (**Figs. 6 and 7**). The mtCa^2+^ uptake rates are slow near the resting [Ca^2+^]_c_ and ER-Mito distance before c-Src activation (Qi, Li et al. 2015), but decreased MAM space may facilitate the increase in basal [Ca^2+^]_m_, followed by *Δψ_m_* depolarization, and increased mROS level (Cao, Adaniya et al. 2019). Under well-coupled conditions, a stimulation of forward flow by itself would be expected to oxidize the electron transport chain (ETC) and thereby lower ROS levels (Nicholls 2004). Acceleration of electron flow within the ETC, however, may also oxidize NADH and NADPH, thereby depleting the anti-oxidative capacity (Aon, Cortassa et al. 2010). Therefore, long-term changes in basal [Ca^2+^]_m_ resulting from decreased MAM space may reduce the amount of antioxidative capacity in the matrix, followed by an increase in ROS levels in mitochondria without a complete loss of *ΔΨ_m_* as we show in **Fig. 6**. Another possibility is that c-Src may have other substrates in the mitochondria and P-Tyr of these substrates could modulate *ΔΨ_m_* independently of the changes in [Ca^2+^]_m_. Although we found the majority of c-Src to be located at the OMM rather than the inside of mitochondria (including their matrix), we could not exclude the possibility that c-Src itself (Miyazaki, Neff et al. 2003), other SFK members (see **Fig. 1**) and/or its downstream tyrosine kinase families (e.g., Pyk2 (O-Uchi, Jhun et al. 2014))) inside the mitochondria also may participate in changing *ΔΨ_m_*, mROS, and cellular respiration. Several mitochondrial matrix proteins have been proposed as specific substrates of c-Src located at the mitochondrial matrix (Miyazaki, Neff et al. 2003, Miyazaki, Tanaka et al. 2006, Ogura, Yamaki et al. 2012, Ogura, Yamaki et al. 2014, Djeungoue-Petga, Lurette et al. 2019, Hebert-Chatelain, Jose et al. 2012), and the phosphorylation of these proteins may modulate mitochondrial respiration. Due to different basal mitochondrial respiration rates and different expression levels of c-Src in the cell types used in these studies, the overall impact of c-Src activation in the matrix on mitochondrial respiration at present is highly controversial. However, several reports (e.g., (Ogura, Yamaki et al. 2014))) showed that overexpression of mitochondria-matrix targeted c-Src does not affect mROS levels. These reports suggest that the increased mROS observed in our experiments is likely not mediated by c-Src activation at the mitochondrial matrix, but more likely from other mitochondrial compartments such as c-Src at the OMM.

Nevertheless, the facilitation of mtCa^2+^ uptake and increased ROS by c-Src could be important components for c-Src-mediated regulation of cell proliferation in various cell types including CFs (Xue, Deng et al. 2018) since 1) recent studies showed the critical correlation between cell proliferation and mtCa^2+^ uptake via mtCUC (Zhao, Li et al. 2019, Koval, Nguyen et al. 2019) and 2) ROS has been established as a fundamental second messenger. Indeed, ROS has been shown as an important cell signaling to activate CF proliferation (Deb, Ubil 2014, Cheng, Cheng et al. 2003, Chen, C. H., Cheng et al. 2006) and we showed that c-Src-mediated P-Tyr of Mfn2 facilitates mtCa^2+^ uptake and ROS generation in primary CFs. Together, these observations suggest that modulating ER-Mio microdomains by c-Src has a significant impact on cellular functions such as cell proliferation.

In conclusion, our results demonstrate a molecular and functional role of c-Src-dependent P-Tyr of Mfn2 in ER-Mito tethering, which is clearly distinct from other PTMs of Mfn2. P-Tyr of Mfn2 may serve as one of the important pathways for the fine-tuning of ER-Mito Ca^2+^ transport under physiological and pathophysiological conditions. Elucidating these unique biophysical characteristics and physiological functions of Mfn2 tyrosine phosphorylation is significant for understanding mitochondrial Ca^2+^ and ROS signaling in the regulation of cellular physiology and pathophysiology.

## Material and methods

### Antibodies, plasmids, and reagents

All antibodies used in this study are shown in **Appendix Table S1**. The plasmids used in this study are listed in **Appendix Table S2**. ER membrane-targeted monomeric yellow fluorescent protein (ER-YFP) was generated from ER membrane-targeted green fluorescent protein (ER-EGFP) (kindly provided by Dr. Tamas Balla, NIH/NICHD, Rockville, MD) (Csordas, Varnai et al. 2010) by replacing the GFP with monomeric YFP (mEYFP) (Zacharias, Violin et al. 2002). All chemicals and reagents were purchased from Sigma-Aldrich (St. Louis, MO) except for: PP2 (Cayman Chemical, Ann Arbor, Michigan); Torin 1 (LC Laboratories, Woburn, MA); thapsigargin (Alomone Labs, Jerusalem, Israel).

### Cell culture and transfection

HEK293T cells (kindly provided by Dr. Keigi Fujiwara (University of Texas MD Anderson Cancer Center, Houston, TX) and IP_3_R1-null HEK293 cells (Kerafast, Inc., Boston, MA) (Alzayady, Wang et al. 2016) were maintained in Dulbecco’s modified Eagle’s medium (DMEM) (HyClone GE Healthcare, Little Chalfont, UK) supplemented with 4.5 g/L glucose, 1 mM sodium pyruvate, 1% L-glutamine, 10% fetal bovine serum (GIBCO, Grand Island, NY, USA), 100 U/mL penicillin, and 100 µg/mL streptomycin (Genesee Scientific, El Cajon, CA) at 37 °C with 5% CO_2_ in a humidified incubator. For maintenance of stable cell lines, antibiotics [1.6 mg/mL of G418 (Corning, Corning, NY) or 0.5 mg/ml puromycin (Gemini Bio-Products, Sacramento, CA)] were added to the medium. The growth medium containing selection antibiotics was refreshed every 48 hr (O-Uchi, Jhun et al. 2013, O-Uchi, Jhun et al. 2014). Transfection was performed using FuGENE HD (Promega, Madison, WI). Twenty-four htr after transfection, cells were detached from the dish using Accutase (Innovative Cell Technologies, San Diego, CA) and re-plated onto glass-bottom dishes (Matsunami USA, Bellingham, WA) or larger size dishes for subsequent live cell imaging, or biochemical experiments after additional 24-48 hr, respectively.

Human adult ventricular cardiac fibroblasts (HCFs) from healthy donors (Cell Applications, San Diego, CA) were maintained in HCF Growth Medium (Cell Applications) at 37 °C with 5% CO_2_ in a humidified incubator (Rizvi, DeFranco et al. 2016). HCFs were infected with adenovirus carrying FLAG (LifeSct, Rockville, MD) or c-Src (Vector Biolabs, Malvern, PA) and used for experiments 48 hr after infection.

For live-cell imaging, all culture medium was replaced by modified Tyrode’s solution containing (136.9 mM NaCl, 5.4 mM KCl, 2 mM CaCl_2_, 0.5 mM MgCl_2_, 0.33 mM NaH_2_PO_4_, 5 mM HEPES, and 5 mM glucose. pH was adjusted to 7.4 using NaOH) (Jhun, O-Uchi et al. 2018).

### Western Blot Analysis

Whole cell lysates from HEK293T cells were prepared with 1x lysis buffer (Cell Signaling Technology, Danvers, MA) containing protease inhibitor cocktail (Roche, Indianapolis, IN or Sigma Aldrich) and 1 mM phenylmethylsulfonyl fluoride (PMSF). Protein concentrations were determined by bicinchoninic acid (BCA) assay. The samples were subjected to SDS-PAGE and transferred to a nitrocellulose membrane for near-infrared fluorescence immunoblotting. Membranes were blocked with blocking buffer (LI-COR Biotechnology, NE and Genesee Scientific, El Cajon, CA) and treated with primary antibodies followed by incubation with fluorescence-conjugated secondary antibodies (LI-COR Biotechnology). Immunoreactive bands were visualized by Odyssey infrared imaging system with Image Studio software (LI-COR Biotechnology). To visualize the levels of phosphorylated MCU and Mfn2, a phosphate-affinity electrophoresis technique (Phos-tag^TM^ SDS-PAGE) was applied using a phosphate-binding molecule (WAKO, Osaka, Japan) (Kinoshita, Kinoshita-Kikuta et al. 2006). Divalent metal ions (Zn^2+^ or Mn^2+^) were added to the SDS-PAGE with Phos-Tag. Protein was blotted onto a nitrocellulose membrane for near-infrared fluorescence immunoblotting.

For Western blots performed by chemiluminescent immunoassay in Fig. 1B, membranes were blocked with 5% fat-free milk or 5% BSA in PBS, probed with a primary antibody, followed by appropriate peroxidase-coupled secondary antibodies. The immunoreactive bands were visualized hrusing SuperSignal West Pico or Femto reagents (Thermo Fisher Scientific, Waltham, MA). Quantitative densitometry was performed in Fiji (Schindelin, Arganda-Carreras et al. 2012) and Image Studio software (LI-COR Biotechnology).

### Immunoprecipitation

Immunoprecipitation (IP) was performed as we previously described (O-Uchi, Jhun et al. 2014, Jhun, O-Uchi et al. 2018). Briefly, whole cell lysates (500 µg) were incubated with primary antibodies (1-2 µg) overnight at 4 °C. Protein A/G Plus Agarose (Santa Cruz) was added to the whole cell lysates for 2 hr at 4 °C and the immunoprecipitate was collected by centrifugation at 1,000 x g for 5 min at 4 °C. After extensively washing the immunocomplexes with lysis buffer, 2x protein sample buffer was added to the immunocomplexes and the beads were dissociated by heating at 95 °C for 5 min. The samples were subjected to Western blotting and immunoreactive bands were visualized by Odyssey infrared imaging system (LI-COR Biotechnology).

### Protein Fractionation

Mitochondria-enriched and cytosolic fractions of HEK293T cells were obtained by protein fractionation using differential centrifugation (O-Uchi, Jhun et al. 2014, O-Uchi, Jhun et al. 2013, Jhun, O-Uchi et al. 2018). Cells were collected from 150-mm dishes, resuspended in isolation buffer (320 mM sucrose, 1 mM EDTA, 10 mM Tris-HCl, pH 7.4) containing protease and phosphatase inhibitor cocktails (Sigma Aldrich). Cells were gently homogenized with a Dounce homogenizer. The mitochondrial and cytosolic proteins containing ER/SR proteins were separated by centrifugation at 17,000 x g for 15 min at 4 °C. The mitochondrial-enriched fraction was re-suspended in lysis buffer containing 1 mM PMSF and protease inhibitor cocktail.

### Mitochondrial Digestion Assay

Mitochondrial digestion was used to determine the sub-mitochondrial localization of CSK as we previously described (O-Uchi, Jhun et al. 2014). The mitochondrial-enriched pellet was re-suspended in 10 mM HEPES buffer (pH 7.2) with 280 mM sucrose and separated into eight samples (30 protein/tube) in 1.7 ml micro-centrifuge tubes. Each tube was treated with 100 µg/ml of proteinase K (Gentrox, Shrewsberry, MA) and differing percentages of digitonin (0.04% to 0.2%) for 15 min at room temperature. In addition to digitonin treatment, triton X-100 (Thermo Fisher Scientific) was used as a positive control to completely lyse the mitochondrial structure so that proteinase K can fully access all proteins. After proteinase K treatment, 10 mM of PMSF was added to each tube to stop the proteinase K digestion and Western blotting was performed. Proteins with known localization were immunoblotted and labeled according to their topology.

### RNA Extraction, Reverse Transcription, and Quantitative Real-Time PCR

Total RNA was isolated from HEK293Tcells using TRIzol™ (Thermo Fisher Scientific). RNA quantity and purity were measured using a NanoDrop 2000 spectrophotometer (NanoDrop Technologies, USA). cDNA was synthesized from 1 µg of total RNA using Superscript™ IV VILO™ Master Mix (Thermo Fisher Scientific). Quantitative real-time PCR (qPCR) was performed using PowerUp SYBR Green Master Mix (Thermo Fisher Scientific) with the Applied Biosystems 7500 Fast Real-Time PCR system (Thermo Fisher Scientific). All other primer sets used for this study were listed in **Appendix Table S3**. The relative transcripts levels were determined by the ΔΔC_T_ method with GAPDH as the endogenous reference gene.

### Quantitative Analyses of Mitochondrial Morphology

Mitochondrial morphology was monitored in cells expressing mitochondrial matrix-targeted DsRed (mt-RFP) (Yoon, Krueger et al. 2003) using a laser scanning confocal microscope Olympus FV3000 (Olympus, Tokyo, Japan) at room temperature (Jhun, O-Uchi et al. 2018, O-Uchi, Jhun et al. 2013). Z-stack images of single HEK293T cells expressing mt-RFP were obtained, and single Z-projection images were created by Fiji software (Schindelin, Arganda-Carreras et al. 2012). Z-projection images were processed through a convolve filter (Jhun, O-Uchi et al. 2018, O-Uchi, Jhun et al. 2013) to obtain isolated and equalized fluorescent pixels. After converting the images, individual mitochondria were subjected to particle analysis to acquire values for circularity and aspect ratio (AR: major axis/minor axis) as previously described (Jhun, O-Uchi et al. 2018, O-Uchi, Jhun et al. 2013). The inverse of circularity was calculated to obtain form factor (FF). Increased AR values indicate long, tubular mitochondria, and increased FF values indicate increased mitochondrial branching and length. A value of 1 for both FF and AR indicates a perfect circle.

### Quantitative Analyses of Mitochondria/ER Membrane Interactions

Interaction between the OMM and ER membranes in live cells was assessed by the Förster resonance energy transfer (FRET) between OMM-targeted CFP (mt-CFP) and ER-mEYFP (**Appendix Table S2**) using confocal microscopy (Olympus FV3000, Olympus) (Jhun, O-Uchi et al. 2018). Briefly, HEK293T cells were co-transfected with an equal amount of mt-CFP and ER-mEYFP. CFP and FRET images were sequentially acquired by the excitation wavelength of 405 nm, and emission wavelength of 475-488 nm and 535-560 nm, respectively. YFP images were obtained as a control (excitation at 488 nm and emission at 530-550 nm).

Interaction between the OMM and ER membranes in fixed cells was assessed by the indirect immunofluorescence for FRET (Konig, Krasteva et al. 2006) by labeling ER membrane SERCA and OMM protein TOM20. Cells were fixed with 4.0 % paraformaldehyde (Thermo Fisher Scientific) at 4 °C for 10 min, permeabilized with saponin at room temperature for 10 min, blocked with 10% goat serum (Cell Signaling Technology), and incubated with primary antibodies overnight, followed by incubation with fluorescent secondary antibodies for 1 hr (O-Uchi, Sasaki et al. 2008). Immunostained images were acquired using an FV3000 laser scanning confocal microscope (Olympus, Tokyo, Japan). Negative control experiments were performed using secondary antibodies without incubation of primary antibodies, which showed no noticeable labeling. Secondary antibodies conjugated with Cyanine3 (Cy3) and Cyanine5 (Cy5) (Thermo Fisher Scientific) were used as a donor and acceptor, respectively. Fluorescent values from non-CMs and CM areas will be quantified using Fiji software (Schindelin, Arganda-Carreras et al. 2012).

### Transmitted Electron Microscopy

ER-Mitochondrial interactions were assessed by transmission electron microscopy (TEM) as we previously reported (O-Uchi, Jhun et al. 2013, Jhun, O-Uchi et al. 2018). HEK293T cells were fixed in 2.5% glutaraldehyde in 0.15 M Na^+^ cacodylate, 2% paraformaldehyde, and 2 mM CaCl_2_ for 2-3 hr at 4 °C, stained with uranyl acetate and lead aspartate, dehydrated in ethanol, and embedded in epoxy resin. Specimens were viewed with a TEM (JEM-1400Plus, JEOL, Tokyo, Japan) at 100 kV (Yamada, Kusakari et al. 2021). Mitochondria size, number, and shape were assessed with Fiji software (Schindelin, Arganda-Carreras et al. 2012).

### Measurement of Cytosolic and Mitochondrial Ca^2+^ Handling

The changes in mitochondrial matrix Ca^2+^ concentration ([Ca^2+^]_mt_) in response to cytosolic Ca^2+^ elevation were measured in cells transfected with a mitochondria-targeted Ca^2+^ biosensor mtRCaMP1h (kindly provided by Dr. Anita Aperia, Karolinska Institutet, Stockholm, Sweden) using a laser scanning confocal microscope Olympus FV3000 (Olympus) at room temperature (Hamilton, Terentyeva et al. 2018). Cells were plated on glass-bottom 35 mm dishes (Matsunami USA) for observation. For the experiments with intact live cells, the cell culture medium was replaced by modified Tyrode’s solution containing 2 mM Ca^2+^ during observation (Hamilton, Terentyeva et al. 2018). The mtCa^2+^ uptake was evoked by IP_3_R-mediated ER Ca^2+^ release induced by applying ATP or angiotensin II to the extracellular solution. For the experiments with permeabilized cells, cells were permeabilized by 50 μM digitonin in a buffer mimicking the cytosolic ionic composition (intracellular buffer: 130 mM KCl, 10 mM NaCl, 2 mM K_2_HPO_4_, 5 mM succinic acid, 5 mM malic acid, 1 mM MgCl_2_, 20 mM HEPES, 1 mM pyruvate, 0.5 mM ATP and 0.1 mM ADP, pH 7) supplemented with 100 μM EGTA (De Stefani, Raffaello et al. 2011). The mtCa^2+^ uptake was evoked by switching the intracellular buffer with 100 μM EGTA to the intracellular buffer with 2 mM EGTA-buffered 3 μM Ca^2+^ (calculated with MAXCHELATOR (Bers, Patton et al. 1994))). The mtRCaMp1h fluorescence was measured with excitation wavelength at 543 nm and emission wavelength at 560-660 nm every 2.5 (for permeabilized cells) or 5 sec (for intact cells). The mtRCaMp1h fluorescence (F) is converted into ΔF/F_0_, which shows the changes in [Ca^2+^]_mt_, where F_0_ stands for initial fluorescence levels.

The changes in cytosolic Ca^2+^ concentration ([Ca^2+^]_cyto_) by IP_3_R-mediated ER Ca^2+^ release were measured in cells loaded with membrane-permeant AM ester form of Fluo-3 (Fluo-3-AM) (Biotium, Fremont, CA, USA) and Fura-Red, Fura 2-TH-AM (Fura-Red-AM) (Setareh Biotech, Eugene, OR, USA) at 37°C and the changes in [Ca^2+^]_cyto_ were measured by confocal microscope Olympus FV3000 (Olympus) at room temperature (O-Uchi, Jhun et al. 2014, Vang, da Silva Goncalves Bos et al. 2021).

### Measurement of Mitochondrial ROS Levels

A mitochondrial matrix-targeted H_2_O_2_-sensitive biosensor, mt-RoGFP2-Orp1 (gift from Dr. Tobias Dick, Deutsches Krebsforschungszentrum, Heidelberg, Germany), was used to measure H_2_O_2_ levels in live HEK293T cells (Meyer, Dick 2010). The fluorescence intensity from mt-RoGFP2-Orp1 (excitation and emission at 510 and 580 nm, respectively) was measured by an epifluorescence microscope equipped with 60x oil TIRF objective lens (Nikon, Tokyo, Japan) and Retiga EXi camera (QImaging, Surrey, BC, Canada) every 10 sec for 6 min (Jhun, O-Uchi et al. 2018). To evaluate the capacity of maximal H_2_O_2_ production in the mitochondrial matrix, tert-butyl hydroperoxide (T-BH, 10 mM) was added to induce oxidative stress on the cells to stimulate maximum mitochondrial H_2_O_2_ production. After cells reached maximal H_2_O_2_ production, 100 mM dithiothreitol (DTT) (Gentrox) was added as a reducing agent to assess the basal mitochondrial H_2_O_2_ level. For measuring mROS in HCFs, cells were stained with mitochondrial superoxide-sensitive dye MitoSOX-Red (Thermo Fisher Scientific) (O-Uchi, Jhun et al. 2013, O-Uchi, Jhun et al. 2014, Jhun, O-Uchi et al. 2018). All data were analyzed using Fiji software (Schindelin, Arganda-Carreras et al. 2012).

### Measurement of mitochondrial membrane potential

Mitochondrial membrane potentials (*Δψ_m_*) were measured by the *Δψ_m_*-sensitive dye TMRE (Thermo Fisher Scientific) in live cells, along with *Δψ_m_-insensitive* dye MitoTracker Deep Red (Thermo Fisher Scientific) (O-Uchi, Jhun et al. 2013). Cells were incubated with 200 mM MitoTracker Deep Red and 500 mM TMRE for 15 minutes at 37 °C to stain the mitochondria and then washed with Tyrode solution. MitoTracker Deep Red fluorescence (excitation and emission at 644 and 665 nm, respectively) and TMRE fluorescence (excitation and emission at 540 and 580 nm, respectively) were simultaneously and time-dependently measured every 10 seconds for 10 minutes with an epifluorescence microscope (Nikon) and Retiga EXi camera (QImaging, Surrey). A mitochondrial oxidative phosphorylation un-coupler carbonyl cyanide m-chlorophenylhydrazone (CCCP, 10 µM) was added to cause *ΔΨ_m_* depolarization. The mitochondrial membrane potential was calculated after the addition of CCCP and normalized to the plateau region for each trace line.

### Mitochondrial respiration measurement

Oxygen consumption rate (OCR) was measured with a Seahorse XFe96 Extracellular Flux analyzer (Agilent Technologies, Santa Clara, CA, USA) as we previously reported (Jhun, O-Uchi et al. 2018). Briefly, HEK293T cells were seeded onto Seahorse XFe96 V3 PS cell culture microplates (Agilent Technologies) at a cell density of 30,000 cells per well and cultured overnight. The hydrated sensor cartridge was loaded in its injection ports with oligomycin (1 μM), carbonyl cyanide-p-trifluoromethoxyphenylhydrazone (FCCP, 0.75 μM), and a mix of rotenone (1 μM) and antimycin A (1 μM), respectively. Seahorse XF DMEM medium (pH 7.4, Agilent Technologies) supplemented with 10 mM glucose, 1 mM pyruvate, and 2 mM L-glutamine was used as assay medium and replaced the culture medium 1 hr before assays. Measurements were taken for three cycles with 3 min mix and 3 min measurement per cycle following each injection. After finishing the assay, cells were stained with Hoechst 33342 (1 μg/ml, Thermo Fisher Scientific) for 30 min at room temperature protected from light, and fluorescent signals were measured with excitation/emission at 350/460 nm using a SynergyMx plate reader (BioTek Instruments, Winooski, VT) for normalization.

## Statistics

All data in the figures are shown as the mean ± standard error (SEM). Unpaired Student’s t-test was performed and a p-value of < 0.05 considered statistically significant. One-way ANOVA followed by Tukey’s post-hoc test was used for multiple comparisons.

## Acknowledgements

The work was supported by American Heart Association (AHA) 13SDG14370008 (to P.Z.), NIH/NIGMS 5P20GM103652 (Targeted project to P.Z.), NIH/NIGMS U54GM115677 (Pilot project to B.S.J.), AHA 18CDA34110091 (to B.S.J), NIH/NHLBI R01HL136757 (to J.O.-U.), NIH/NIGMS P30GM114750 (Pilot project to J.O.-U.), NIH/NIGMS P30GM110759 (Pilot project to J.O.-U.), AHA 16SDG27260248 (to J.O.-U.), American Physiological Society (APS) 2017 Shih-Chun Wang Young Investigator Award (to J.O.-U.). The authors thank Ms. Michelle King, Ms. Amy K Landi, Dr. Xiaofei Li, Dr. Nedyalka Valkov, Ms. Gayathri Dileepan, and Ms. Dongqin Yang for their technical assistance.

## Author contributions

P.Z. and J.O.-U. conceived and designed research; P.Z., K.F., J.H.S., Y,.S. M.L., Ja.M., I.C., I.P., M.K., B.N., T.T., Y.K., M.W.C., S.M.A., Jy.M., B.S.J., and J.O.-U. performed experiments; P.Z., K.F., J.H.S., Y.S., M.L., Ja.M., M.W.C., K.D., B.S.J., and J.O.-U. analyzed data; P.Z, K.F., T.T., Y.K., M.W.C., Ja.M., U.M., B.S.J., and J.O.-U. interpreted results of experiments; P.Z., K.F., J.H.S., Y.S., B.S.J. and J.O.-U. prepared figures; J.O.-U. drafted manuscript; P.Z., Ja.M., T.T., M.W.C., U.M., B.S.J., and J.O.-U. edited and revised manuscript; all authors approved final version of manuscript.

## Conflict of interest

No conflicts of interest, financial or otherwise, are declared by the author(s).

## Notes

### Competing Interest Statement

The authors have declared no competing interest.

### Summary of Updates

All the text and data sets

