## Supplementary material for "c-Src-dependent phosphorylation of Mfn2 regulates endoplasmic reticulum-mitochondria tethering": Fig. EV

### Expanded View Figure and Figure Legends

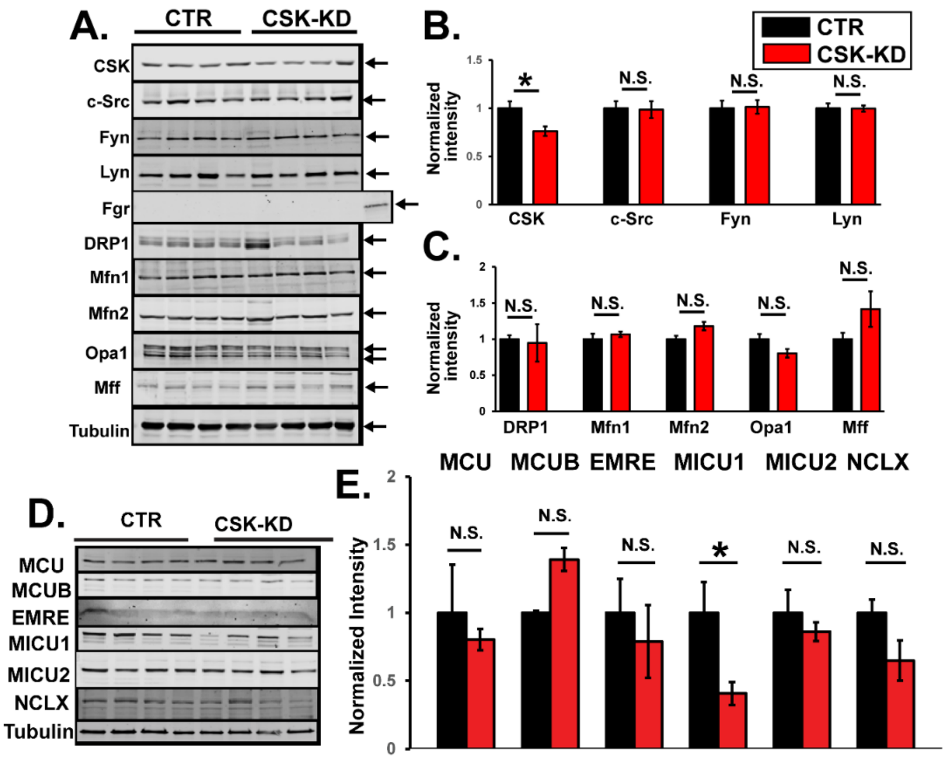


**Fig. EV1. Expression levels of Src family, mitochondrial fission/fusion proteins, and mtCa$^{2+}$ handling proteins in HEK293T cell stably knockdown CSK**

**A.** Representative immunoblots of whole cell lysates obtained from different passages of HEK293T cells stably overexpressing PLKO.1 (control, CTR) or CSK- shRNA (CSK-KD). The sample from each lane is from different passages. A lysate of HEK293T cells overexpressing Fgr was used as a positive control for Fgr. Mfn1, Mitofusin 1; Mff, mitochondrial fission factor; Opa1, optic atrophy-1; DRP1, dynamin-related protein 1. **B.** Summary data of SFK expression levels. Each band intensity was normalized by tubulin and expressed relative to CTR (*n*=4 for each group). **C.** Summary data of mitochondrial fission/fusion protein expression levels. Each band intensity was normalized by tubulin and expressed relative to CTR (*n*=4 for each group). **D.** Protein expression levels of mtCa$^{2+}$ handling proteins in whole cell lysates obtained different passages of CTR and CSK-KD HEK293T cells. **E.** Summary data of D. Each band intensity was normalized by tubulin and expressed relative to CTR (*n*=4 for each group).

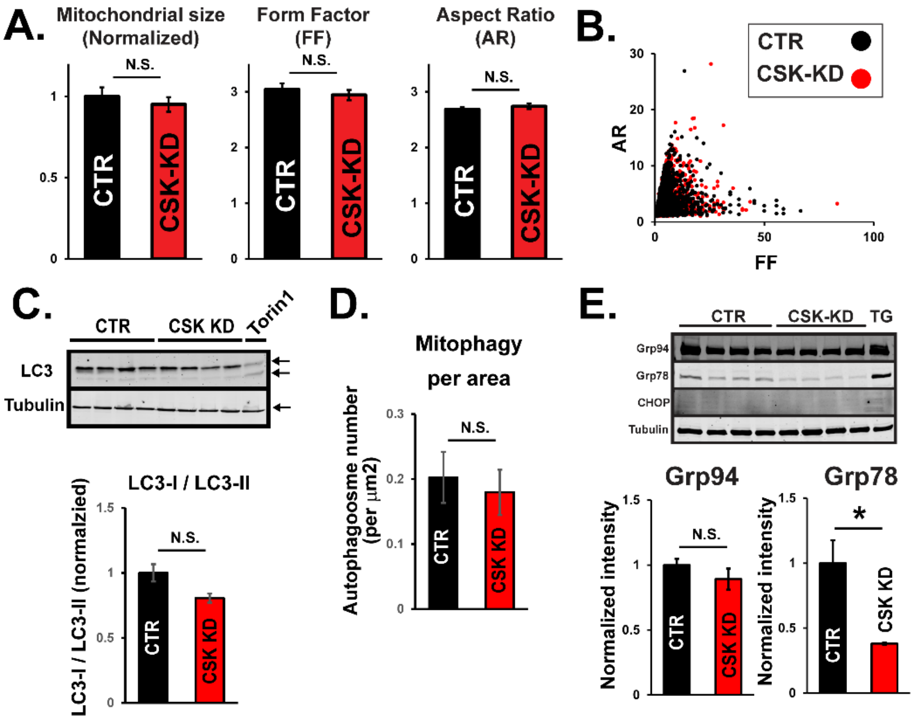


**Fig. EV2**. **CSK-knockdown does not enhance mitochondrial fusion, mitophagy, or ER stress signaling.**

**A.** Summary data of mitochondrial size, form factor (FF), and aspect ratio (AR) in CTR and CSK-KD HEK293T cells (*n* = 4249 and 4394, respectively). Cells were transfected with matrix-targeted DsRed (mt-RFP) and analyzed with live cell imaging using confocal microscopy. N.S., not significant. **B.** Comparison of AR and FF of individual mitochondria in CSK-KD cells vs. CTR cells. **C.** *(Top)*: Representative immunoblots of LC3-I/LC3-II obtained from CTR and CSK-KD HEK293T cells. The sample from HEK293T cells treated with Torin1 is shown as a positive control that changes LC3-I/LC3-II ratio. *(Bottom):* Summary data for LC3-I/LC3-II ratio. The value of LC3-I/LC3-II ratio from CSK-KD cells was expressed relative to CTR (*n*=4). **D.** Mitophagosome number counted from TEM images of CTR and CSK-KD HEK293T cells (*n*=60, and 83, respectively) (see also Fig. 2). **E.** *(Top):* Representative immunoblots of ER stress markers, Grp94, Grp78, and CHOP obtained from CTR and CSK-KD HEK293T cells. The sample from HEK293T cells treated with thapsigargin (TG, 100 nM for overnight) is shown as a positive control that increases ER stress. *(Bottom):* Summary data (*n*=4). Grp94 and Grp78 band intensities were normalized to tubulin and expressed relative to CTR. * $p<0.05$.

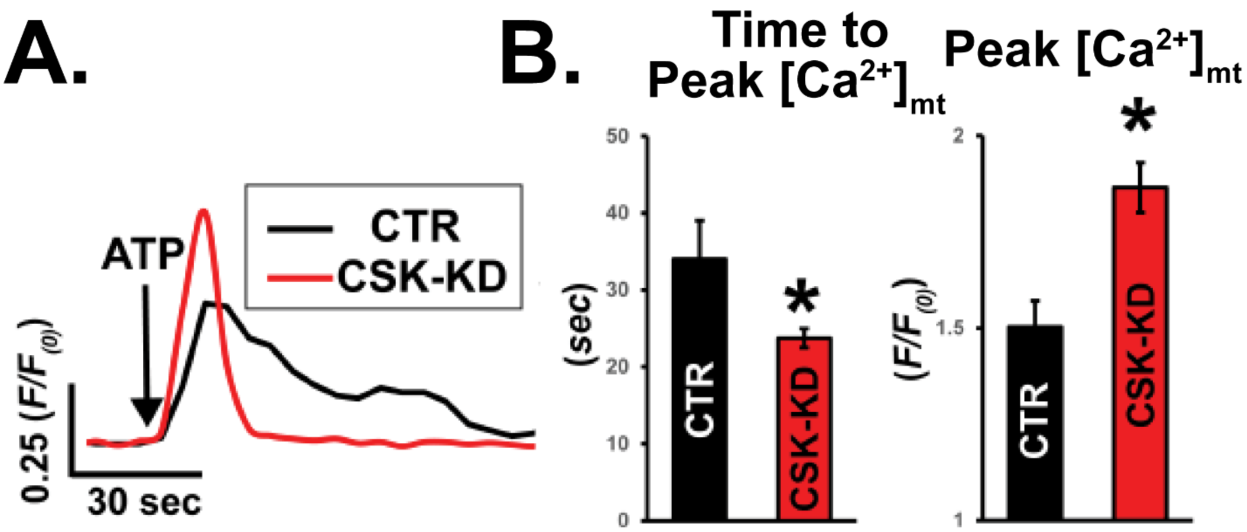


**Fig. EV3**. **CSK-knockdown increases mtCa$^{2+}$ uptake in response to cytosolic Ca$^{2+}$ elevation**

**A.** Representative traces of changes in Ca$^{2+}$ concentration in mitochondrial matrix ([Ca$^{2+}$]$_m$) obtained from individual HEK293T cells stably overexpressing PLKO.1 (control: CTR) or CSK-shRNA (CSK knockdown: CSK-KD) in response to $G_{\alpha q/11}$ protein-coupled P2Y receptor stimulation by 1 mM ATP. [Ca$^{2+}$]$_m$ was assessed with the mitochondrial matrix-targeted Ca$^{2+}$ biosensor mt-RCamp1h. **B.** Summary data of A. Black and red bars are from CTR and CSK-KD cells, respectively. ($n$ = 36 and $n$ = 56 for CTR and CSK-KD cells, respectively). *$P$ <0.05.

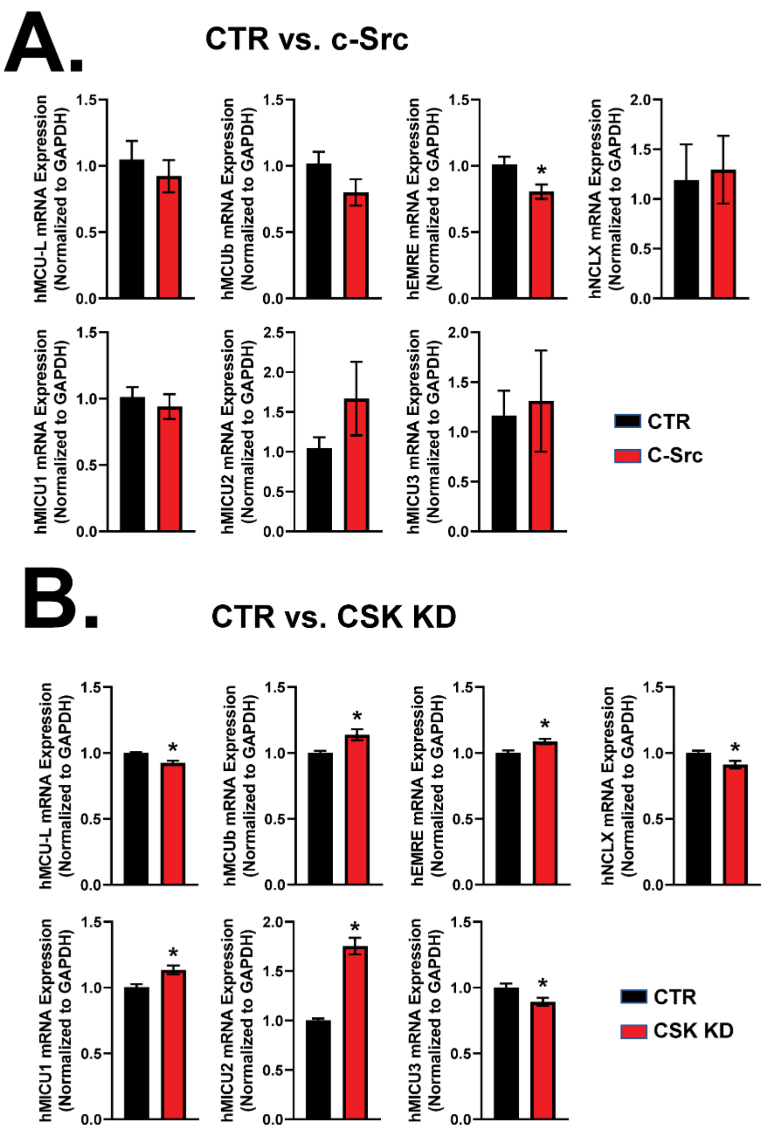


**Fig. EV4. Effect of c-Src overexpression or CSK-KD on mtCa$^{2+}$ handling proteins in HEK293T cells.**

**A.** Real-time qPCR analysis of major mtCa$^{2+}$ handling proteins in HEK293T cells stably overexpressing c-Src compared to the cells expressing pcDNA3.1(+) empty vector (CTR) ($n$ = 6 for each group). *$P$ <0.05. The mRNA level of each gene was normalized to that in GAPDH and expressed relative to CTR. **B.** Real-time qPCR analysis of major mtCa$^{2+}$ handling proteins in HEK293T cells stably overexpressing shRNA targeting CSK (CSK-KD) compared to the cells expressing pLKO.1 empty vector (CTR). ($n$ = 6 for each group). *$P$ <0.05.
