## Appendix for "c-Src-dependent phosphorylation of Mfn2 regulates endoplasmic reticulum-mitochondria tethering"

**Table of contents**

**Fig. S1. SFK-specific inhibitor PP2 concentration-dependently inhibits on SFK activity in CSK-KD and control cells**

**Fig. S2. CSK-knockdown increases the amount of $IP_3$ receptor in the MAM**

**Fig. S3. Alignment of amino acid sequence of Mfn2 from different species**

**Table S1. List of commercial primary antibodies used in this study**

**Table S2. List of plasmids used**

**Table S3. List of primers used**

**Reference list for the Appendix**

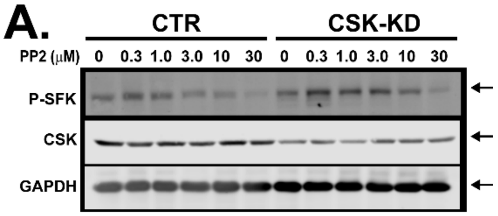


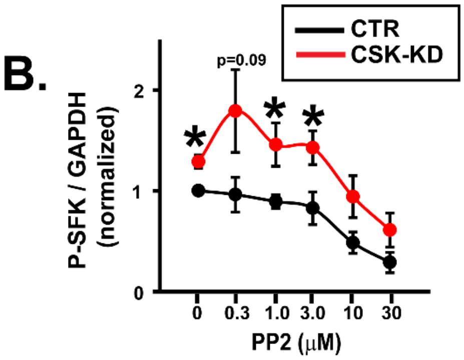


**Fig. S1. SFK-specific inhibitor PP2 concentration-dependently inhibits on SFK activity in CSK-KD and control cells**

**A.** Representative near-infrared fluorescence immunoblots of whole cell lysates obtained from HEK293T cells stably overexpressing PLKO.1 (control, CTR) or CSK- shRNA (CSK-KD) that were treated with the SFK inhibitor PP2 at indicated concentrations for 30 min. To assess the activity of SFKs, total SFK autophosphorylation levels were detected by a specific antibody raised against a synthetic phosphopeptide corresponding to residues surrounding $Tyr^{419}$ of human c-Src, which can cross-react with the autophosphorylation sites of all other SFKs (Du, Wang et al. 2020). **B.** Summary data of panel A (CSK-KD in red, CTR in black). Band intensity for P-SFK was normalized by GAPDH and expressed relative to CTR without PP2 (0 μM) ($n$ = 4 for each group). The values of Y axis are expressed as log scale. *$P$ <0.05, compared to CTR at each PP2 concentration.

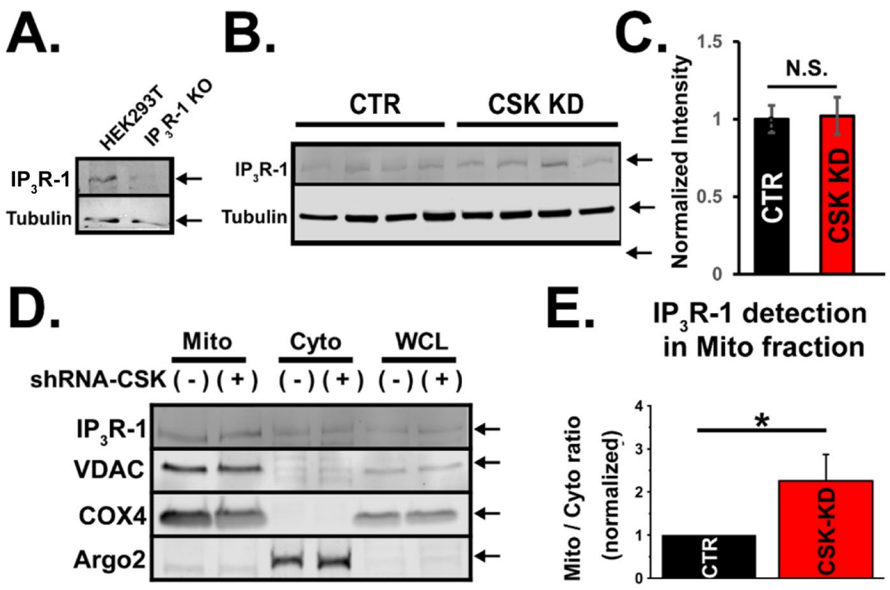


**Fig. S2. CSK-knockdown increases the amount of $IP_3$ receptor in the MAM**

**A.** Verification of the specificity of a commercial antibody against $IP_3$ receptor isoform 1 ($IP_3$R-1) using whole cell lysates from HEK293T cells and an $IP_3$R-1-knockout HEK293 cell line by a specific antibody against rat $IP_3$R-1. **B.** Immunoblots of $IP_3$R-1 in whole cell lysates obtained from HEK293T cells stably overexpressing CSK-shRNA (CSK-KD) or PLKO.1 (control; CTR). **C.** Summary data of B. $IP_3$R-1 band intensities were normalized by tubulin and expressed relative to CTR ($n$ = 4). N.S., not significant. **D.** Immunoblots of protein fractionations from CSK-KD and CTR cells. $IP_3$R-1 was used to assess the MAM protein amount in the mitochondrial fraction. Mito, mitochondrial fraction; Cyto, cytosolic fraction including ER and plasma membrane; WCL, whole cell lysate. VDAC, COX4, and Argo2 were detected as markers for outer mitochondrial membrane, mitochondrial matrix, and cytosol, respectively. **E.** Summary data of the $IP_3$R-1 expression ratio between Mito and Cyto ($n$ = 4). $IP_3$R-1 expression levels in Mito and Cyto in each cell line were normalized by the amount of COX4 and Argo2, respectively, before calculating the Mito/Cyto ratio. The values were expressed relative to CTR. *$P$ <0.05.

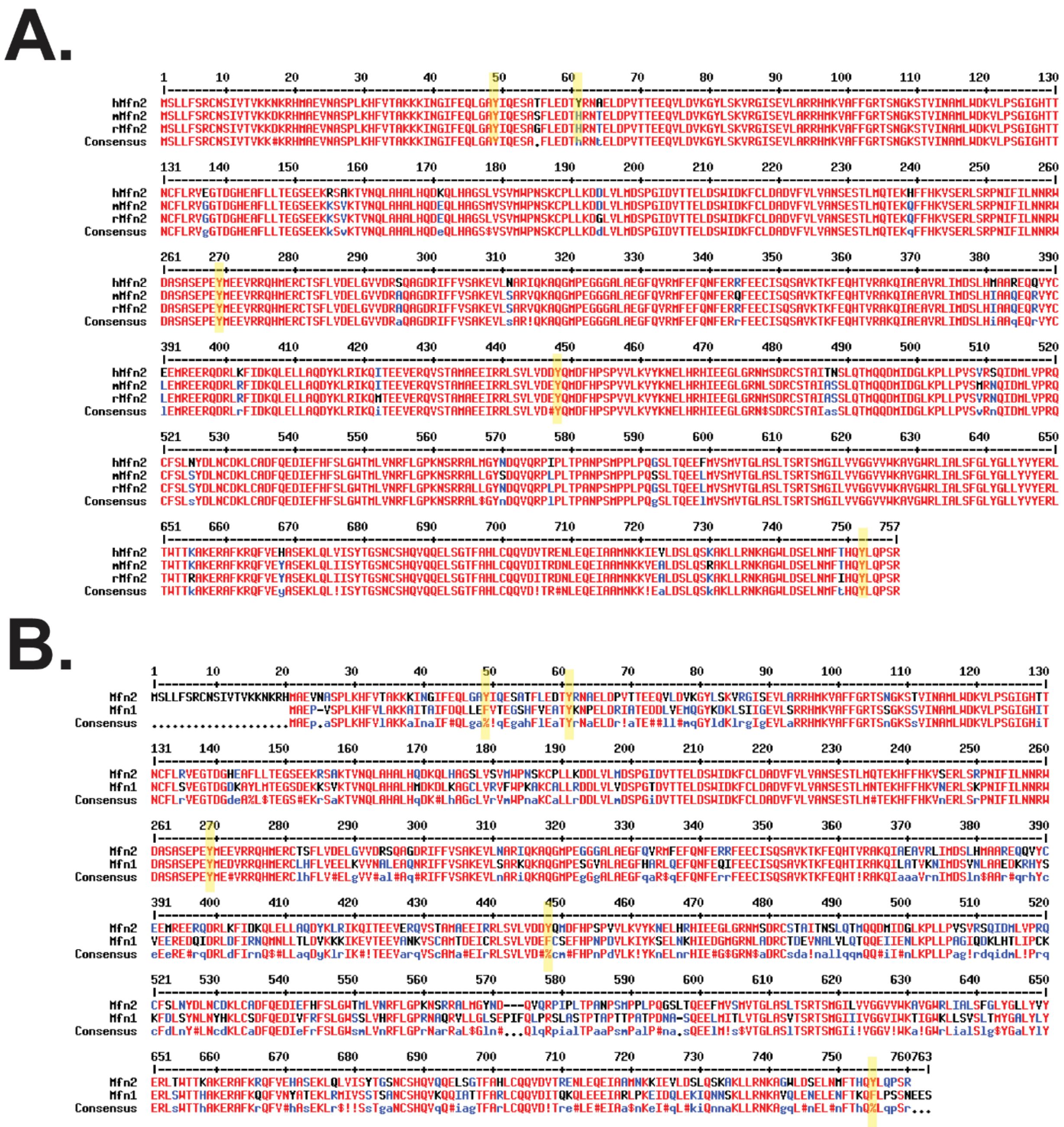

**Fig. S3. Comparison of amino acid sequences of Mfn2 from different species and between human Mfn2 and Mfn1**

**A.** Sequence alignment of human, mouse, and rat Mfn2. Multiple sequence alignment were performed by MultAlin (http://multalin.toulouse.inra.fr/multalin/multalin.html). Yellow highlights show the locations of the candidate Tyr residues for c-Src-specific phosphorylation sites in human Mfn2. **B.** Sequence alignment of human Mfn2 and Mfn1.

**Table S1. List of commercial primary antibodies used in this study**

| Targeted protein | Type | Company | Catalog number | Immunogen |
|---|---|---|---|---|
| Mitofusin 1 (Mfn1) | Rabbit polyclonal | Abcam, Cambridge, United Kingdom | ab104274 | Synthetic peptide corresponding to human Mfn1 1 aa 683-732 |
| Mitochondrial fission factor (MFF) | Rabbit polyclonal | Sigma-Aldrich, St. Louis, MO | ABS2087 | His-tagged full length human recombinant mitochondrial fission factor (Mff). |
| Phospho-Src family-Tyr416 | Rabbit monoclonal | Cell Signaling Technology, Danvers, MA | 6943S | Synthetic phosphopeptide corresponding to residues surrounding Tyr419 of human Src protein. |
| c-Src | Mouse monoclonal | Cell Signaling Technology | 2110 | Recombinant fusion protein containing an N-terminal fragment of human Src |
| c-Src | Rabbit monoclonal | Cell Signaling Technology | 2123 | Recombinant fusion protein corresponding to residues 1-110 of human Src protein |
| Fyn | Rabbit polyclonal | Cell Signaling Technology | 4023T | Synthetic peptide corresponding to residues surrounding Ser25 of human Fyn |
| Lyn | Rabbit monoclonal | Cell Signaling Technology | 2796 | Synthetic peptide corresponding to the amino-terminal sequence of human Lyn |
| Fgr | Rabbit polyclonal | Cell Signaling Technology | 2755S | Synthetic peptide corresponding to residues close to the amino terminus of human Fgr |
| CSK | Rabbit monoclonal | Cell Signaling Technology | 4980S | Synthetic peptide corresponding to residues surrounding amino acid residue Val399 of human CSK. |
| Argonaute 2 (Argo2) | Rabbit monoclonal | Cell Signaling Technology | 2897S | Synthetic peptide corresponding to mouse Argonaute 2 |
| Voltage-dependent anion channel | Rabbit polyclonal | Cell Signaling Technology | 4866S | Synthetic peptide corresponding to the |

| | | | | |
|---|---|---|---|---|
| (VDAC) | | | | amino terminus of human VDAC-1 |
| Cytochrome c oxidase subunit 4 (COX4) | Rabbit monoclonal | Cell Signaling Technology | 4850 | Synthetic peptide corresponding to residues surrounding Lys29 of human COX IV |
| Glyceraldehyde 3-phosphate dehydrogenase (GAPDH) | Rabbit monoclonal | Cell Signaling Technology | 2118 | Synthetic peptide near the carboxy terminus of human GAPDH |
| HTRA2 (Omi) | Rabbit monoclonal | Cell Signaling Technology | 9745S | Synthetic peptide corresponding to residues surrounding Phe341 of human HtrA2/Omi protein |
| Glucose-regulated protein 94 (Grp94) | Rabbit monoclonal | Cell Signaling Technology | 20292S | Synthetic peptide corresponding to residues surrounding Leu397 of human Grp94 protein |
| Glucose-regulated protein 78 (Bip/Grp78) | Rabbit monoclonal | Cell Signaling Technology | 3177S | Synthetic peptide corresponding to residues surrounding Gly584 of human BiP |
| C/EBP-homologous protein (CHOP) | Mouse monoclonal | Cell Signaling Technology | 2895S | Synthetic peptide corresponding to the sequence of human CHOP |
| human influenza hemagglutinin (HA) | Mouse monoclonal | Sigma-Aldrich | H9658 | Synthetic peptide corresponding to a fragment of HA, conjugated to KLH |
| HA | Rabbit polyclonal | Sigma-Aldrich | H6908 | Synthetic peptide corresponding to amino acid residues of HA, conjugated to KLH |
| Tubulin | Mouse monoclonal | Sigma-Aldrich | T5168 | Sarkosyl-resistant filaments from *S. purpuratus* (sea urchin) sperm axonemes |
| Mfn2 | Rabbit polyclonal | Sigma-Aldrich | M6319 | Synthetic peptide corresponding to amino acid residues 38-55 of human Mfn2 with C-terminal added cysteine, conjugated to KLH |
| Phospho-tyrosine | Mouse | Sigma-Aldrich | 05-321 | Phosphotyramine-KLH |

| | | | | |
|---|---|---|---|---|
| | monoclonal | | | |
| Optic atrophy-1 (OPA1) | Mouse monoclonal | BD Bioscience, Franklin Lakes, NJ | 12607 | Human OPA1 aa 708-830 |
| Dynamin-related protein 1 (DRP1) | Mouse monoclonal | Cell Signaling Technology | 14647S | recombinant protein specific to the amino terminus of human DRP1 protein |
| Cyclophilin D (Cyp-D) | Mouse monoclonal | Thermo Fisher Scientific, Waltham, MA | 455900 | Recombit rat Cyclophilin F |
| LC3A/B | Rabbit monoclonal | Cell Signaling Technology | 12741 | Synthetic peptide corresponding to residues surrounding Leu44 of human LC3B protein (conserved in LC3A) |
| Inositol 1,4,5-trisphosphate receptor ($IP_3R$) receptor isoform 1 ($IP_3R1$) | Rabbit polyclonal | Alomone Labs, Jerusalem Israel | ACC-019 | Peptide (C)RIGLLGHPPHMNVNPQQPA, corresponding to amino acid residues 2732-2750 of rat IP3R1 (Accession P29994). Intracellular, C-terminus |
| Myc | Mouse monoclonal | BioLegend | 626802 | Amino acids 408-439, C-terminal region of human c-Myc |
| MCU | Rabbit polyclonal | Sigma-Aldrich | HPA016480 | N-terminal region of human MCU (HHRTVHQRIASWQNLGAVYCSTVVPSDDVTVVYQNGLPVISVRLPSRRERCQFTLKPISDSVGVFLRQLQEEDRGIDRVAIYSPDGVRVAASTGIDLLLLDDFKLV). |
| SERCA1/2/3 | Mouse monoclonal | Santa Cruz Biotechnology | sc-271669 | Amino acids 1-300 mapping at the N-terminus of SERCA1 of human origin |
| MCUb | Rabbit polyclonal | Proteintech, Rosemont, IL | 20387-1-AP | MLQRGLWPWRTRLLPTPGTWRPARPWPLPPPPQVLRVKLCGNVKYYQSHHYSTVVPPDEITVIYRHGLPLVTLTLPSRKERCQFVVKPMLSTVGSFLQDLQNEDKGIKTAAIFTADGNMISASTLMDILLMNDFKLVINKIAYDVQCPKREKPSNEHTAEMEHMKSLVHRLFTILHLEESQKKREHHLLEKIDHLK |

| | | | | |
|---|---|---|---|---|
| | | | | EQLQPLEQVKAGIEAHSEAKTSGLLWAGLALLSI |
| MICU1 | Rabbit polyclonal | Sigma-Aldrich | HPA037480 | LKGKLTIKNFLEFQRKLQHDVLKLEFERHDPVDGRITERQFGGMLLAYSGVQSKKLTAMQRQLKKHFKEGKGLTFQEVENFFTFL |
| MICU2 | Rabbit polyclonal | Abcam | ab101465 | Recombinant fragment corresponding to human MICU2 aa 95-238 |
| EMRE | Rabbit polyclonal | Santa Cruz Biotechnology | sc-86337 | Peptide mapping at the C-terminus of human C22orf32 |
| NCLX | Rabbit polyclonal | Proteintech | 21430-1-AP | YVVTVILCTWIYQRQRRGSLFCPMPVTPEILSDSEEDRVSSNTNSYDYGDEYRPLFFYQETTAQILVRALNPLDYMKWRRKSAYWKALKVFKLP |
| TOM20 | Rabbit polyclonal | Proteintech | 11802-1-AP | MVGRNSAIAAGVCGALFIGYCIYFDRKRRSDPNFKNRLRERRKKQKLAKERAGLSKLPDLKDAEAVQKFFLEEIQLGEELLAQGEYEKGVDHLTNAIAVCGQPQQLLQVLQQTLPPPVFQMLLTKLPTISQRIVSAQSLAEDDVE |

### Table S2. List of plasmids used

| Inserted gene | Vector Backbone | Source/Provider | Company | Notes | Ref. |
|---|---|---|---|---|---|
| Mitochondrial matrix-targeted GFP (mt-GFP) | pEGFP | Dr. Yisang Yoon, Augusta University, Augusta, GA | | | (O-Uchi, Jhun et al. 2013) |
| Mitochondrial matrix-targeted DsRed (mt-REP) | pDsRed | Dr. Yisang Yoon | | | (O-Uchi, Jhun et al. 2013) |
| Mitochondrial matrix-targeted $Ca^{2+}$ biosensor (mt-RCaMP1h) | pDsRed2 | Dr. Anita Aperia, Karolinska Institutet, Stockholm, Sweden | | | (Hamilton, Terentyeva et al. 2018) |
| HA-tagged human Mfn2 | pcDNA3.1(+) | Dr. Takumi Koshiba, Kyushu University, Fukuoka, Japan | | | (Yasukawa, Oshiumi et al. 2009) |
| OMM-targeted monomeric ECFP (mt-CFP) | pcDNA3 | | | | (Jhun, O-Uchi et al. 2018) |
| ER-surface targeted EGFP containing the C-terminal segment of Sac1 (ER-EGFP) | pEGFP-C1 | Dr. Tamas Balla, NIH/NICHD, Rockville, MD | | | (Csordas, Varnai et al. 2010) |
| Mitochondrial $H_2O_2$-sensitive biosensor, mt-RoGFP2-Orp1 | pLPCX | Dr. Tobias Dick, FF Heidelberg, Germany | Addgene, Watertown, MA | Addgene plasmid # 64992; http://n2t.net/addgene:64992; RRID:Addgene_64992 | (Meyer, Dick 2010) |
| empty | pLKO.1-puro | Dr. Bob Weinberg, Whitehead Institute for Biomedical research, Cambridge, MA | Addgene | Addgene plasmid # 8453; http://n2t.net/addgene:8453; RRID:Addgene_8453 | |
| Mouse c-Src-WT | pCMV5 | Drs. Joan Brugge & Peter Howley, Harvard Medical School, Boston, MA | Addgene | Addgene plasmid # 13663; http://n2t.net/addgene:13663 | |
| Mouse c-Src K295R Y527F (c-Src-DN) | pCMV5 | Drs. Joan Brugge & Peter Howley | Addgene | Addgene plasmid # 13657; http://n2t.net/addgene:13657; RRID:Addgene_13657 | |

| | | | | | |
|---|---|---|---|---|---|
| Flag-tagged human Fyn | pcDNA3 | Dr. Lars Rönnstrand, Lund University, Lund, Sweden | Addgene | Addgene plasmid # 74509 ; http://n2t.net/addgene:74509 ; RRID:Addgene_74509 | (Chougule, Kazi et al. 2016) |
| empty | pWZL-Neo-Myr-Flag-DEST | Dr. Jean Zhao, Dana-Farber Cancer Institute, Boston MA | | Addgene plasmid # 15300 http://n2t.net/addgene:15300; RRID: Addgene_15300 | (Boehm, Zhao et al. 2007) |
| V5-tagged human Lyn | pLX304 | | GeneCopoeia, Rockville, MD | | |
| Human Fgr | pReceiver-M02 | | GeneCopoeia | | |
| N/A | pEGFP-C1 | | Promega, Madison, WI | | |
| shRNA targeted to coding region of human CSK mRNA (CSK shRNA) | | RNAi Consortium Broad Institute of MIT and Harvard University, Cambridge MA | Sigma Aldrich | CCGGCGAGGAGGTGTACTTTGAGAACTCGAGTTCTCAAAGTACACCTCCTCGTTTTT | |
| shRNA targeted to 3'-UTR of human Mfn2 (Mfn2 shRNA) | | RNAi Consortium Broad Institute of MIT and Harvard University, Cambridge MA | Sigma Aldrich | CCGGGCTCAGTGCTTCATCCCATTTCTCGAGAAATGGGATGAAGCACTGAGCTTTTTG | |
| ER membrane-targeted monomeric YFP (ER-mYFP) | pEGFP-C1 | | | generated from ER-EGFP | |

### Table S3. List of primers used

| Gene Name | NCBI Gene ID | FWD Sequence (5'-3') | REV Sequence (5'-3') |
|---|---|---|---|
| GAPDH | 2597 | 5'-TCG GAG TCA ACG GAT TTG-3' | 5'-CAA CAA TAT CCA CTT TAC CAG AG-3′ |
| MCU | 90550 | 5'-ATATTCCTGGGACATCATGGAG-3' | 5'-GGATAAACATATTCCTGGCGTG-3′ |
| MCUb | 55013 | 5′-GGC CTT CCC TTG GTA ACA CT-3' | 5'-GTT GCC ATC TGC TGT GAA GA-3′ |
| MICU1 | 10367 | 5'-GAG GCA GCT CAA GAA GCA CT-3' | 5'-CAA ACA CCA CAT CAC ACA CG-3’ |
| MICU2 | 221154 | 5'-GGC AGT TTT ACA GTC TCC GC-3' | 5'-AAG AGG AAG TCT CGT GGT GTC-3’ |
| MICU3 | 286097 | 5’-AGA TGA ATT TAA ACG TGC CG-3’ | 5’-GGA GTC TGT CTT TCA TAA TTC C-3’ |
| EMRE | 91689 | 5'-TGT CGG GAC ACT CAT TAG CA-3' | 5'-GCT GAT AGG GAA GGC AGA GA-3’ |
| NCLX | 80024 | 5'-GCC AGC ATT TGT GTC CAT TT-3' | 5'-AAT TCG TCT CGG CCA CTT AC-3' |
